# TENT5A anchors STING at the endoplasmic reticulum to block SEC24B-dependent trafficking and suppress cGAS-STING signaling

**DOI:** 10.64898/2026.07.24.739811

**Authors:** Shengnan Zheng, Pei-hui Wang

## Abstract

The cGAS-STING pathway is a central sensor of cytosolic DNA that drives type I interferon responses during infection and tumor surveillance. Activation of STING requires coordinated trafficking from the endoplasmic reticulum (ER) to the Golgi, yet how this spatial transition is regulated by ER-resident factors remains incompletely understood. Here we identify TENT5A as an ER-localized negative regulator of STING signaling. Genetic depletion of TENT5A markedly enhances cGAS-STING-dependent type I interferon production in response to cytosolic DNA and DNA virus infection. Conversely, TENT5A overexpression suppresses downstream signaling and antiviral gene expression. Mechanistically, TENT5A directly interacts with the C-terminal tail of STING and competes with the COPII adaptor SEC24B, thereby preventing STING incorporation into COPII vesicles and blocking its ER-to-Golgi trafficking. This ER retention inhibits STING oligomerization, TBK1 recruitment and IRF3 activation, ultimately attenuating type I interferon responses. In vivo, myeloid-specific deletion of TENT5A enhances systemic interferon responses, restricts herpesvirus replication, and improves host survival. Moreover, TENT5A deficiency promotes CD8⁺ T cell effector activation within the tumor microenvironment and suppresses tumor growth in a melanoma model. Together, our findings define TENT5A as an ER checkpoint that spatially controls STING trafficking through competition with COPII machinery, revealing vesicular sorting as a regulatory layer of innate immune activation.

## Introduction

Cytosolic DNA sensing through the cGAS-STING pathway is a central mechanism by which mammalian cells detect infection and malignant transformation. Activation of this pathway initiates the production of cyclic GMP–AMP (cGAMP), which binds and activates STING, triggering TBK1-dependent phosphorylation of IRF3 and subsequent type I interferon (IFN-I) responses that are essential for antiviral immunity and tumor immune surveillance^[1–3]^. Given its potent immunostimulatory capacity, STING activity is subject to multilayered regulatory control to prevent aberrant inflammation and autoimmunity^[4]^.

A defining yet underappreciated feature of STING signaling is its strict dependence on intracellular trafficking. Unlike canonical cytosolic signaling cascades, STING activation is initiated at the endoplasmic reticulum (ER) but requires rapid translocation to ER–Golgi intermediate compartments and the Golgi apparatus, where signaling complexes are assembled and sustained^[5]^. This spatial transition is not merely permissive but is a prerequisite for downstream signaling output, placing membrane trafficking at the core of innate immune signal activation^[5]^. Accordingly, defects in STING trafficking lead either to immunodeficiency or to autoinflammatory disease, underscoring the need for precise spatial control of this pathway^[6]^.

Recent studies have begun to connect STING trafficking to components of the secretory pathway, particularly COPII vesicle formation machinery^[7]^. The ER exit of STING requires the COPII adaptor SEC24, suggesting that canonical vesicle sorting machinery directly participates in immune signaling regulation^[8]^. However, how STING selectively engages COPII components in response to cytosolic DNA, and whether this process is actively restrained by ER-resident regulatory factors, remains largely unknown. In particular, it is unclear whether dedicated ER-localized “checkpoint” proteins exist that gate STING access to the secretory trafficking machinery prior to ER export.

Here, we identify TENT5A^[9]^ as an ER-resident regulatory factor that functions as a spatial checkpoint for STING activation. We demonstrate that TENT5A directly binds the C-terminal regulatory region of STING and competitively inhibits its interaction with the COPII adaptor SEC24B. This competition prevents STING incorporation into COPII vesicles, thereby blocking ER-to-Golgi translocation and downstream signaling complex assembly. As a consequence, TENT5A restrains STING oligomerization, TBK1 recruitment, and IRF3 activation, resulting in suppression of type I interferon responses.

Importantly, this ER checkpoint mechanism has functional consequences in vivo. Genetic ablation of TENT5A enhances type I interferon responses during DNA virus infection, improves viral clearance, and strengthens host survival. Moreover, TENT5A deficiency potentiates antitumor immunity by promoting CD8⁺ T cell effector activation and suppressing tumor growth in vivo. Together, our findings define TENT5A as a previously unrecognized ER checkpoint that governs STING activation by regulating access to COPII-dependent trafficking machinery, revealing a spatial licensing mechanism that integrates vesicular transport with innate immune signaling.

## Results

### TENT5A negatively regulates cGAS-STING-induced IFN-I signaling

To elaborate the regulatory network of the cGAS-STING pathway, we first identified candidate proteins (Gene01–Gene22) with potential binding capacity to cGAS or STING via protein interaction databases and constructed their overexpression plasmids. A dual-luciferase reporter assay for IFN-β was used for preliminary functional screening in HEK293T cells stably expressing STING. The results revealed that TENT5A specifically suppresses cGAS-STING-triggered IFN-β responses (Supplementary Figure 1 A). Gradient overexpression of TENT5A further demonstrated that its inhibitory effect on the cGAS-STING pathway increases in a dose-dependent manner (Supplementary Figure 1 B), implying that TENT5A acts as a negative regulator of cGAS-STING signaling activity.

To further characterize TENT5A regulatory function, we generated HeLa and THP-1 cell lines stably expressing TENT5A-3×Flag (TENT5A) or the empty vector (E.V.) and confirmed robust overexpression at both the mRNA level (Supplementary Figure 4A) and the protein level (Supplementary Figure 4F, G, H), as described below.

We first examined the effect of TENT5A overexpression on IFN-I signaling at the transcriptional level. Stimulation of HeLa-TENT5A cells with CT-DNA for 6 h and 9 h significantly reduced the mRNA levels of *IFNB1*, *CXCL10*, AND *ISG56* compared to E.V. controls (Supplementary Figure 4B). Consistent results were obtained following HSV-1 infectionat the same time points (Supplementary Figure 4C), confirming that TENT5A suppresses IFN-I induction at the transcriptional level in response to both synthetic DNA and live DNA virus. At the protein level, transient co-expression of TENT5A with cGAS and STING in HEK293T cells markedly reduced cGAS-STING-induced phosphorylation of TBK1 and IRF3 (Supplementary Figure 4D). Furthermore, gradient transfection of TENT5A expression plasmid into HeLa cells (the plasmid itself acts as a dsDNA stimulus) resulted in dose-dependent reductions in STING, TBK1, and IRF3 phosphorylation (Supplementary Figure 4E), confirming that TENT5A specifically and dose-dependently inhibits cGAS-STING pathway activation.To further consolidate these findings in stable cell-line models, we stimulated HeLa-TENT5A and HeLa-E.V. cells with CT-DNA or infected them with HSV-1 and assessed signaling by western blot. Under CT-DNA stimulation, TENT5A-overexpressing HeLa cells exhibited significantly reduced ISG15 protein expression and markedly attenuated phosphorylation of STING, TBK1, and IRF3, with no change in total protein levels (Supplementary Figure 4F). Identical findings were obtained following HSV-1 infection (Supplementary Figure 4G). To verify that these observations extend to a human monocytic cell line, we repeated the CT-DNA stimulation experiment in THP-1 cells stably expressing TENT5A or E.V.; the results were fully concordant with those in HeLa cells, with suppressed ISG15 expression and reduced phosphorylation of STING, TBK1, and IRF3 in TENT5A-overexpressing cells (Supplementary Figure 4H). Together, these overexpression data establish that TENT5A is a potent negative regulator of cGAS-STING-induced IFN-I signaling across multiple human cell types.

To complement the overexpression studies with loss-of-function evidence, we generated THP-1 cell lines stably expressing two independent shRNAs targeting TENT5A (shTENT5A-1 and shTENT5A-2) or a non-targeting control (shGFP). Efficient knockdown was confirmed at the mRNA level by qPCR (Supplementary Figure 5A) and at the protein level by western blot (Supplementary Figure 5D, 5E). Following CT-DNA stimulation, TENT5A knockdown cells showed significantly elevated mRNA levels of *IFNB1*, *CXCL10*, and *ISG56* compared to shGFP controls at multiple doses and time points (Supplementary Figure 5B), with concordant increases in STING, TBK1, and IRF3 phosphorylation as well as ISG15 protein expression by western blot (Supplementary Figure 5D). Following HSV-1 infection, TENT5A knockdown similarly enhanced *IFNB1, CXCL10, and ISG56* mRNA induction while reducing *HSV-1-TK* mRNA levels (Supplementary Figure 5C), and augmented the phosphorylation of STING, TBK1, and IRF3, upregulated ISG15, and decreased the viral immediate-early protein ICP0 (Supplementary Figure 5E). These results from two independent shRNA lines provide robust genetic evidence that endogenous TENT5A restrains cGAS-STING-mediated IFN-I responses in human monocytic cells.

To further exclude potential off-target effects of shRNA-mediated knockdown, we employed CRISPR/Cas9 to generate THP-1 cells in which TENT5A was stably deleted using two independent sgRNAs (sgTENT5A-1 and sgTENT5A-2), with sgCtrl as the negative control. Successful knockout was confirmed by western blot demonstrating near-complete loss of TENT5A protein (Supplementary Figure 6B, 6C). Phenotypically, sgTENT5A cells recapitulated the knockdown phenotype: CT-DNA stimulation induced significantly higher *IFNB1, CXCL10, and ISG56* mRNA levels in both sgTENT5A lines relative to sgCtrl (Supplementary Figure 6A, left), along with enhanced phosphorylation of STING, TBK1, and IRF3 and upregulated ISG15 (Supplementary Figure 6B). HSV-1 infection likewise elicited augmented antiviral gene expression and reduced *HSV-1-TK* mRNA (Supplementary Figure 6A, right), with correspondingly increased pathway phosphorylation, elevated ISG15, and reduced ICP0 protein (Supplementary Figure 6C). The concordance between independent knockdown and knockout approaches strongly supports the conclusion that TENT5A is an endogenous suppressor of cGAS-STING-induced IFN-I production.

To validate the regulatory function of TENT5A in a physiologically relevant primary cell model, we generated myeloid-specific conditional knockout mice (*Tent5a*^fl/fl^ Lyz2-Cre) by crossing *Tent5a*^fl/fl^ mice with Lyz2-Cre mice, and isolated bone marrow-derived macrophages (BMDMs) from littermate *Tent5a*^fl/fl^ and *Tent5a*^fl/fl^ Lyz2-Cre mice. Complete ablation of TENT5A protein was confirmed by western blot (Figure 1C, 1D, 1E). Stimulation of *Tent5a*^fl/fl^ Lyz2-Cre BMDMs with CT-DNA or infection with HSV-1 induced significantly higher mRNA levels of *Ifnb1*, *Cxcl10*, *Isg56*, while *HSV-1-TK* mRNA was markedly reduced, compared to *Tent5a*^fl/fl^ controls (Figure 1A). IFN-β protein secretion measured by ELISA in culture supernatants was also significantly elevated in *Tent5a*^fl/fl^ Lyz2-Cre BMDMs following CT-DNA stimulation, HSV-1 infection, or VACV infection (Figure 1B). To confirm that these effects were robust across stimulation doses, we further tested VACV70 at 1 μg/mL and 2 μg/mL, which yielded consistent enhancement of *Ifnb1*, *Cxcl10*, and *Isg56* mRNA in TENT5A-deficient BMDMs (Supplementary Figure 2A). Likewise, infection with HSV-1 at a higher MOI of 2 confirmed the dose-independent nature of the enhancement (Supplementary Figure 2B). At the protein level, western blot analysis showed that TENT5A deficiency significantly augmented CT-DNA-induced phosphorylation of STING, IRF3, and TBK1 (Figure 1C), HSV-1-induced phosphorylation of these same molecules alongside reduced ICP0 expression (Figure 1D), and VACV-induced STING-TBK1-IRF3 axis activation (Figure 1E), collectively demonstrating that TENT5A suppresses cGAS-STING signal transduction in primary macrophages in response to multiple DNA stimuli.

**Figure 1.**
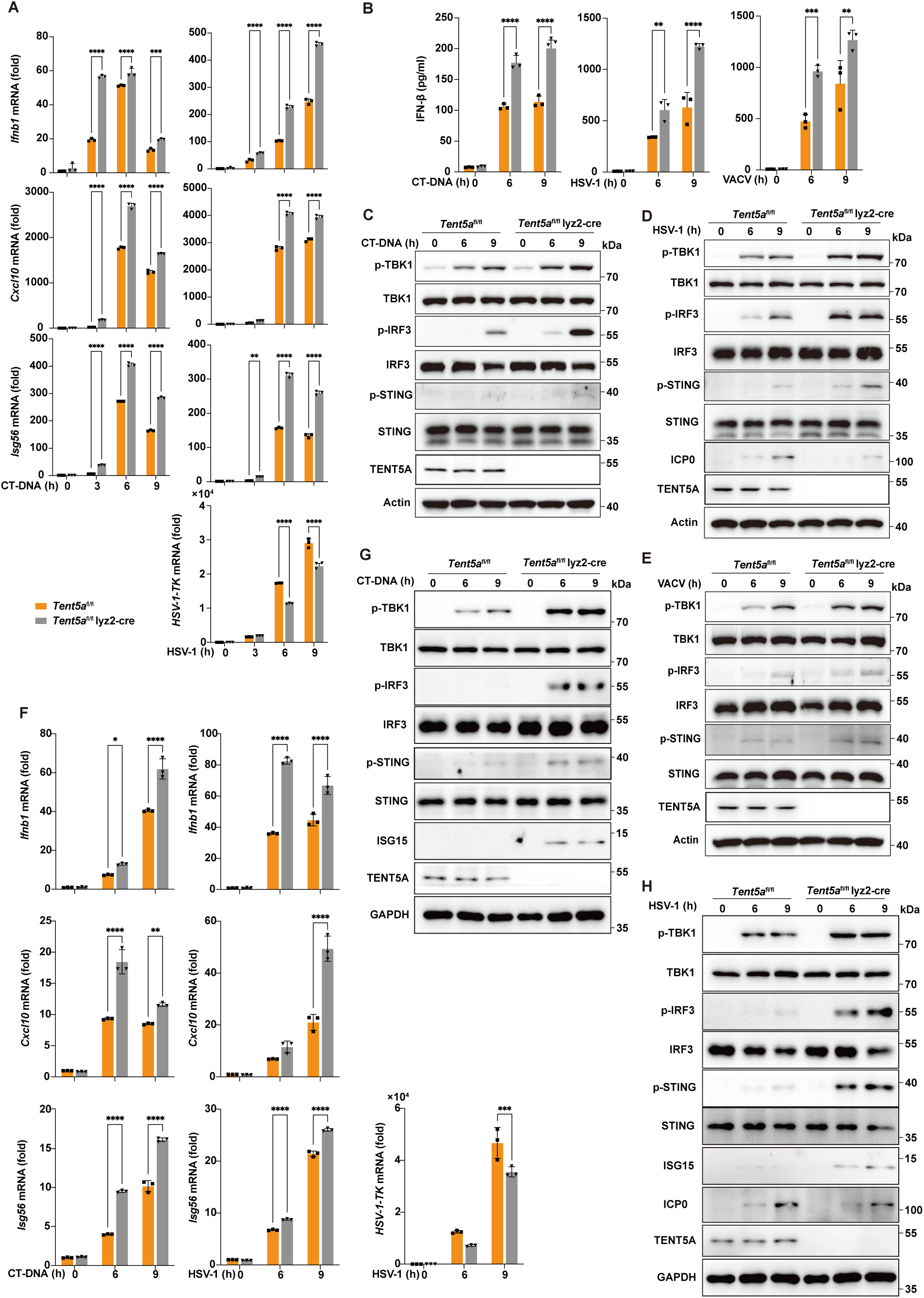
TENT5A negatively regulates cGAS-STING-induced IFN-I signaling. (A) qPCR analysis of *Ifnb1, Cxcl10, Isg56,* and *HSV-1-TK* mRNA in *Tent5a*^fl/fl^ and *Tent5a*^fl/fl^ Lyz2-Cre BMDMs following stimulation with CT-DNA (1 μg/mL) or infection with HSV-1 (MOI = 1) for the indicated time points. (B) ELISA analysis of IFN-β in culture supernatants from *Tent5a*^fl/fl^ and *Tent5a*^fl/fl^ Lyz2-Cre BMDMs stimulated with CT-DNA (1 μg/mL), infected with HSV-1 (MOI = 1), or infected with VACV (MOI = 1) for the indicated time points. (C) Western blot analysis of phosphorylated and total STING, IRF3, and TBK1 in lysates of *Tent5a*^fl/fl^ and *Tent5a*^fl/fl^ Lyz2-Cre BMDMs stimulated with CT-DNA (1 μg/mL) for the indicated time points. (D) Western blot analysis of ICP0 and phosphorylated and total STING, IRF3, and TBK1 in lysates of *Tent5a*^fl/fl^ and *Tent5a*^fl/fl^ Lyz2-Cre BMDMs infected with HSV-1 (MOI = 1) for the indicated time points. (E) Western blot analysis of phosphorylated and total STING, IRF3, and TBK1 in lysates of *Tent5a*^fl/fl^ and *Tent5a*^fl/fl^ Lyz2-Cre BMDMs infected with VACV (MOI = 1) for the indicated time points. (F) qPCR analysis of *Ifnb1, Cxcl10, Isg56,* and *HSV-1-TK* mRNA in *Tent5a*^fl/fl^ and *Tent5a*^fl/fl^ Lyz2-Cre PMs following by stimulation with CT-DNA (1 μg/mL) or infection with HSV-1 (MOI = 1) for the indicated time points. (G) Western blot analysis of ISG15 and phosphorylated and total STING, IRF3, and TBK1 in lysates of *Tent5a*^fl/fl^ and *Tent5a*^fl/fl^ Lyz2-Cre PMs stimulated with CT-DNA (1 μg/mL) for the indicated time points. (H) Western blot analysis of ICP0, ISG15, and phosphorylated and total STING, IRF3, and TBK1 in lysates of *Tent5a*^fl/fl^ and *Tent5a*^fl/fl^ Lyz2-Cre PMs infected with HSV-1 (MOI = 1) for the indicated time points. Data are presented as mean ± S.D. Two-way ANOVA; \**P <* 0.05, \*\**P <* 0.01, \*\*\**P <* 0.001, \*\*\*\**P <* 0.0001 (A, B, F).

To confirm that the regulatory phenotype of TENT5A is not restricted to BMDMs, we extended our analysis to peritoneal macrophages (PMs) isolated from littermate *Tent5a*^fl/fl^ and *Tent5a*^fl/fl^ Lyz2-Cre mice. Western blot confirmed complete loss of TENT5A protein in *Tent5a*^fl/fl^ Lyz2-Cre PMs (Figure 1G, 1H). Following CT-DNA stimulation or HSV-1 infection, *Tent5a*^fl/fl^ Lyz2-Cre PMs displayed markedly elevated mRNA levels of *Ifnb1*, *Cxcl10*, and *Isg56*, accompanied by reduced *HSV-1-TK* mRNA (Figure 1F). At the protein level, CT-DNA stimulation produced enhanced phosphorylation of STING, TBK1, and IRF3 together with upregulated ISG15 in TENT5A-deficient PMs (Figure 1G), while HSV-1 infection additionally resulted in decreased ICP0 expression alongside the same augmented signaling profile (Figure 1H). The consistent findings across BMDMs and PMs establish that TENT5A is a broadly acting negative regulator of cGAS-STING-mediated IFN-I signaling in primary myeloid cells.

To determine whether the inhibitory function of TENT5A extends beyond myeloid cells to non-immune cell types, we examined *Tent5a*^+/+^ and *Tent5a*^-/-^ mouse embryonic fibroblasts (MEFs). Knockout was confirmed at the mRNA level by qPCR (Supplementary Figure 3A) and at the protein level by western blot (Supplementary Figure 3D, 3E). *Tent5a*^-/-^ MEFs stimulated with VACV70 or CT-DNA exhibited significantly increased *Ifnb1*, *Cxcl10*, and *Isg56* mRNA expression compared to *Tent5a*^+/+^ controls (Supplementary Figure 3B). VACV infection further validated the enhanced antiviral response in *Tent5a*^-/-^ MEFs, evidenced by elevated *Ifnb1*, *Cxcl10*, and *Isg56* mRNA and reduced viral replication marker *VACV-J2* mRNA (Supplementary Figure 3C). Western blot analysis confirmed that CT-DNA-induced phosphorylation of STING, TBK1, and IRF3 and ISG15 upregulation were all significantly enhanced in *Tent5a*^-/-^ MEFs (Supplementary Figure 3D), as were HSV-1-induced signaling responses, with concomitant reduction of ICP0 protein (Supplementary Figure 3E). These results demonstrate that the cell-type-independent suppression of cGAS-STING signaling by TENT5A is a conserved function, operating in both immune and non-immune cell contexts.

### TENT5A targets STING

Having established that TENT5A is a negative regulator of cGAS-STING-induced IFN-I signaling in multiple cell models, we next sought to identify the specific node within the pathway at which TENT5A exerts its inhibitory function. To this end, we performed epistasis experiments by co-transfecting TENT5A together with individual pathway components spanning successive tiers of the signaling cascade including cGAS, STING, TBK1, IKKε, and a constitutively active IRF3 mutant (IRF3-5D) into HEK293T cells and measuring transcriptional output by dual-luciferase reporter assays. TENT5A significantly suppressed IFN-β-Luc, IFN-λ1-Luc, and ISRE-Luc reporter activity driven by cGAS+STING co-expression or by STING alone, but had no appreciable effect on reporter activity induced by TBK1, IKKε, or IRF3-5D overexpression (Figure 2B), placing the inhibitory action of TENT5A at or upstream of TBK1. To corroborate these findings at the endogenous gene expression level, we measured *IFNB1*, *CXCL10*, *ISG54*, and *IFNL1* mRNA by qPCR under the same co-transfection conditions. Consistent with the reporter data, TENT5A selectively suppressed the transcriptional induction of all four genes driven by STING alone or by cGAS+STING, while leaving TBK1-, IKKε-, and IRF3-5D-induced gene expression unaffected (Figure 2A). The convergent results from two independent readouts, reporter assays and endogenous mRNA quantification, unambiguously place the action of TENT5A upstream of TBK1.

**Figure 2.**
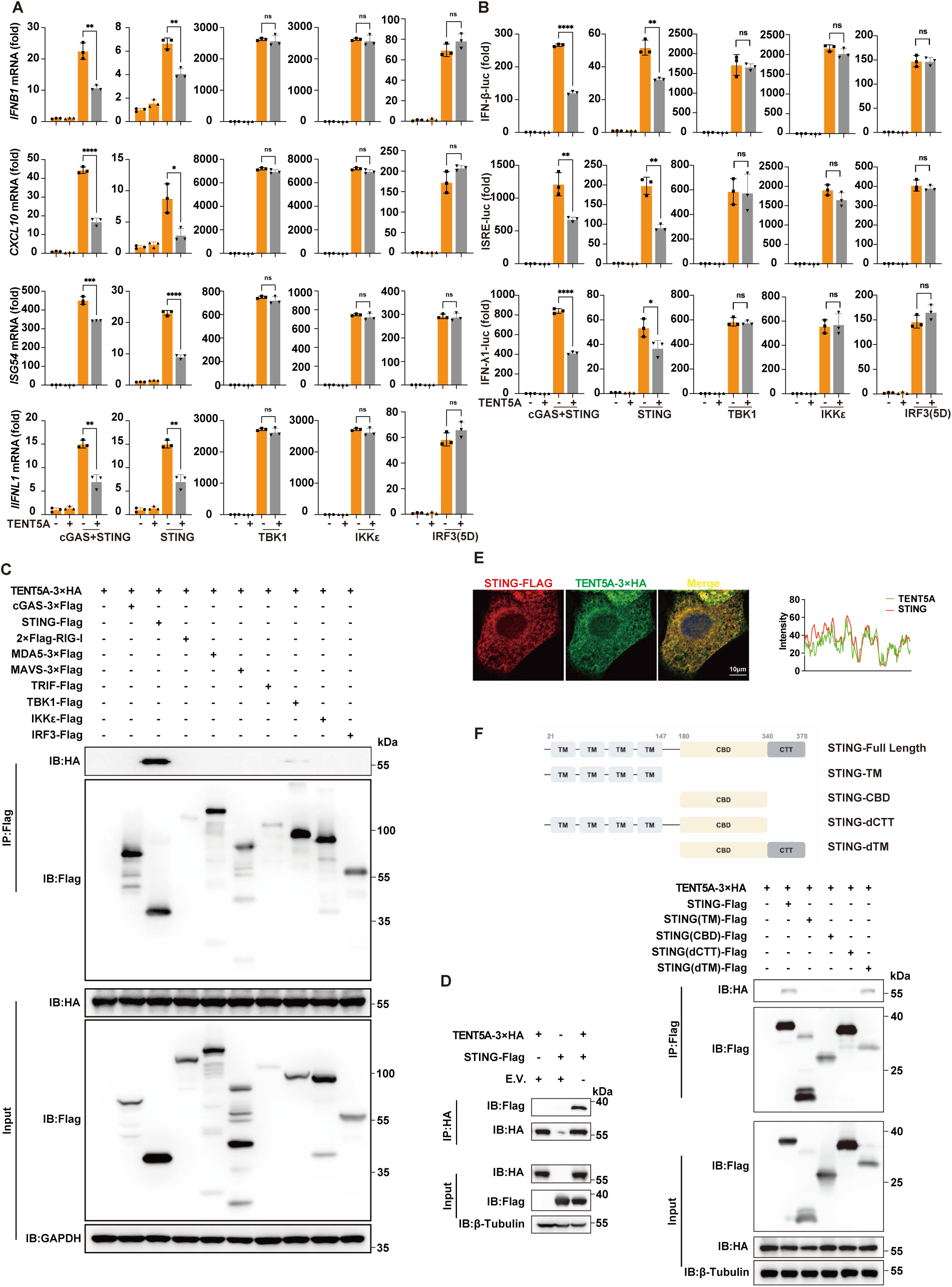
TENT5A targets STING. (A) qPCR analysis of *IFNB1, CXCL10, ISG54,* and *IFNL1* mRNA in HEK293T cells co-transfected with the indicated expression plasmids. (B) Dual-luciferase reporter assays for IFN-β-Luc, IFN-λ1-Luc, and ISRE-Luc reporters in HEK293T cells co-transfected with the indicated expression and reporter plasmids; pRL-TK served as the internal control. (C) Co-IP was performed of TENT5A with key molecules of the innate nucleic acid-sensing pathway. Lysates from HEK293T cells expressing TENT5A-3×HA and the indicated constructs were immunoprecipitated with anti-Flag magnetic beads. (D) Co-IP analysis of the interaction between TENT5A and STING. Lysates from HEK293T cells expressing TENT5A-3×HA and STING-Flag were immunoprecipitated with anti-HA agarose beads. (E) Colocalization analysis of TENT5A-3×HA and STING-Flag. Colocalization was quantified using ImageJ. Scale bar, 10 μm. (F) Mapping the STING domain required for TENT5A binding. HEK293T cells were co-transfected with TENT5A-3×HA and full-length or truncated STING-Flag constructs. Lysates were immunoprecipitated with anti-Flag magnetic beads. A schematic of STING truncations is shown above. Data are presented as mean ± S.D. Two-tailed unpaired Student’s t-test; \**P <* 0.05, \*\**P <* 0.01, \*\*\**P <* 0.001, \*\*\*\**P <* 0.0001 (A, B).

Given that the epistasis data converged on upstream of TBK1 as the functional target of TENT5A, we next aimed to pinpoint the exact molecular target underlying the inhibitory function of TENT5A. We performed a Co-IP screen in HEK293T cells by co-expressing TENT5A-3×HA with Flag-tagged versions of key innate nucleic acid-sensing pathway molecules, encompassing the core cGAS-STING components (cGAS, STING), the RIG-I/MDA5-MAVS axis (RIG-I, MDA5, MAVS), the TLR adaptor TRIF, and shared downstream effectors (TBK1, IRF3, IKKε). Immunoprecipitation with anti-Flag magnetic beads revealed that TENT5A specifically co-precipitated with STING and with none of the other pathway components tested (Figure 2C), establishing that the TENT5A–STING interaction is highly selective and not a general feature of innate immune signaling complexes. To further validate this interaction, we performed a reciprocal Co-IP experiment in HEK293T cells co-expressing TENT5A-3×HA and STING-Flag, using anti-HA agarose beads for immunoprecipitation. STING-Flag was efficiently co-immunoprecipitated with TENT5A-3×HA (Figure 2D), confirming a bona fide and bidirectional physical association between the two proteins.

Physical interaction requires spatial proximity within the cell. We therefore examined the subcellular localization of TENT5A to determine whether it occupies a compartment compatible with STING engagement. Immunofluorescence analysis in HeLa cells expressing TENT5A-3×HA showed extensive co-localization of TENT5A with the ER-Tracker, but minimal overlap with the mitochondrial-Tracker or the Golgi-Tracker (Supplementary Figure 7A), demonstrating that TENT5A is predominantly an ER-resident protein — the same compartment where STING resides in its resting, inactive state. Consistent with this, co-expression of TENT5A-3×HA and STING-Flag in HeLa cells revealed prominent co-localization of the two proteins within the cytoplasm, with a high degree of fluorescence signal overlap (Figure 2E), providing direct spatial evidence that TENT5A and STING occupy the same intracellular compartment. To further explore whether this association persists into the activated state of STING, we assessed the co-localization of exogenous TENT5A-3×HA with endogenous phosphorylated STING (p-STING) in HeLa cells. We found that TENT5A exhibited no obvious co-localization with p-STING (Supplementary Figure 7B). These data indicate that TENT5A predominantly associates with inactive STING rather than phosphorylated, activated STING, suggesting that TENT5A dissociates from STING following its phosphorylation and subsequent activation.

To define the structural basis of the TENT5A–STING interaction, we performed domain-mapping experiments using a series of STING truncation mutants. Full-length STING and STING truncation mutants were co-expressed with TENT5A-3×HA in HEK293T cells, and Co-IP was performed using anti-Flag magnetic beads. TENT5A co-immunoprecipitated with full-length STING and with all truncation mutants that retained the C-terminal tail (CTT) domain of STING, whereas constructs lacking the CTT failed to co-precipitate TENT5A (Figure 2F). These results identify the CTT of STING as the domain required for TENT5A binding. Notably, the CTT is also the critical regulatory domain that mediates TBK1 recruitment and IRF3 phosphorylation following STING activation, raising the possibility that TENT5A and downstream effectors may compete for CTT occupancy. Collectively, the data in this section establish that TENT5A specifically targets STING as its direct binding partner within the cGAS-STING pathway, interacts with STING through the CTT domain at the ER, and does not engage other innate immune signaling molecules, providing a molecular foundation for the mechanistic studies described below.

### TENT5A inhibits multiple key activation events of STING

Having established that TENT5A specifically targets STING, we next sought to determine at which step of the STING activation cascade TENT5A exerts its inhibitory function. STING activation proceeds through a series of sequential events: at steady state, STING resides at the endoplasmic reticulum (ER) membrane as an inactive dimer^[10]^; binding of cGAMP to STING triggers conformational rearrangement of STING, thereby promoting its transport from the ER to the ER-Golgi intermediate compartment (ERGIC) and Golgi^[11]^. While trafficking along this secretory pathway, STING forms functional oligomers and recruits the downstream effector kinase TBK1 as well as the transcription factor IRF3^[12]^. TBK1 further phosphorylates IRF3, leading to IRF3 dimerization and nuclear translocation^[13]^. Consequently, the transcription of type I interferons (IFN-I) is robustly activated. We therefore systematically investigated the effect of TENT5A at each of these key nodes. Since STING activation can be negatively regulated through ubiquitin-mediated proteasomal degradation, we first tested whether TENT5A affects STING protein stability^[14, 15]^. Western blot analysis of HEK293T cells co-transfected with STING-Flag and increasing amounts of TENT5A-3×HA showed that TENT5A overexpression had no appreciable effect on total STING protein levels (Figure 3A), ruling out the possibility that TENT5A promotes STING degradation and indicating that its regulatory mechanism operates downstream of protein stability.

**Figure 3.**
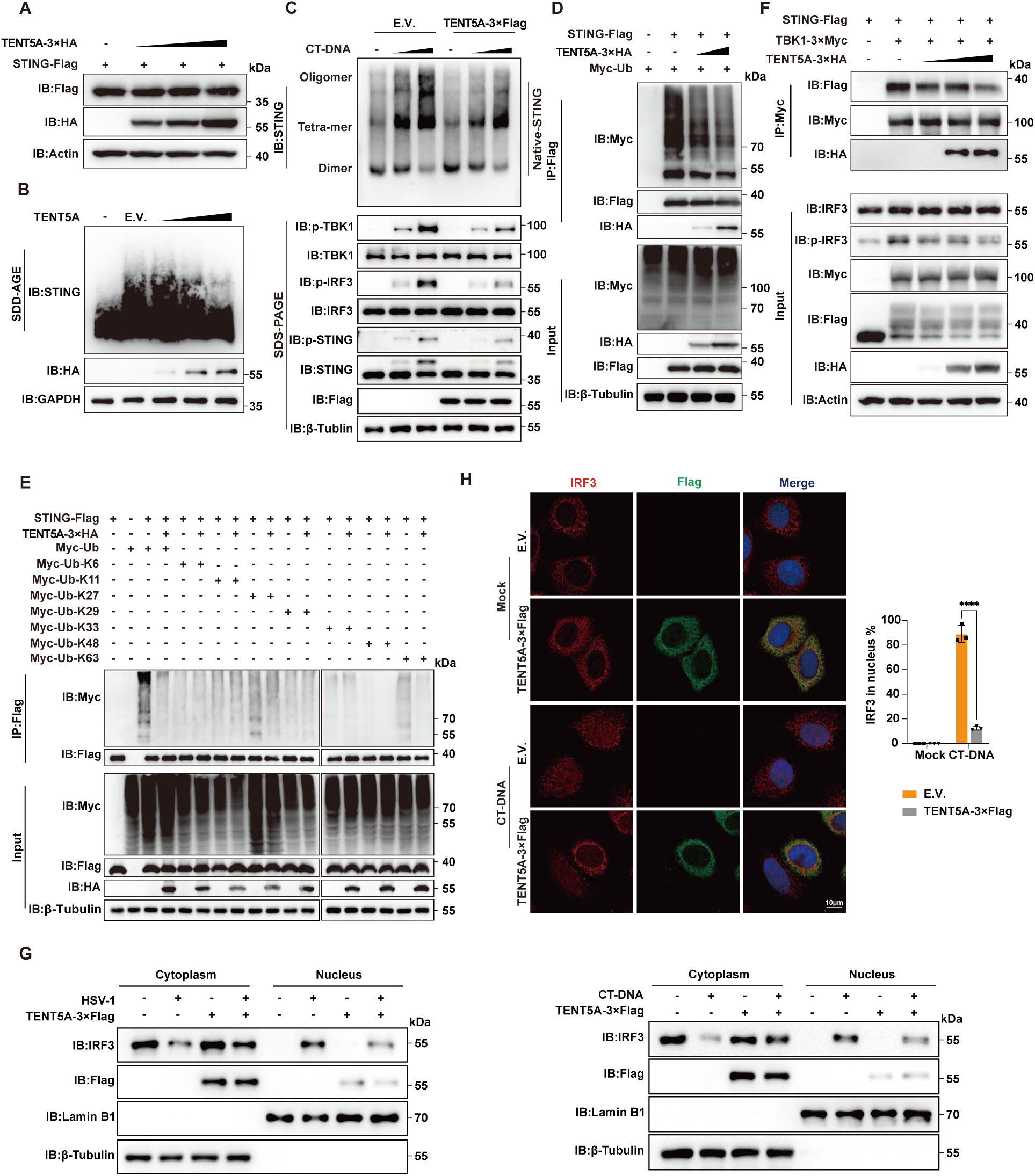
TENT5A inhibits multiple key activation events of STING. (A) Western blot analysis of STING protein stability in HEK293T cells co-transfected with STING-Flag and increasing amounts of TENT5A-3×HA plasmids. (B) SDD-AGE analysis of endogenous STING oligomerization in HeLa cells transfected with E.V. or increasing dose of TENT5A-3×HA plasmids. (C) Native PAGE and SDS-PAGE analysis of STING dimerization, oligomerization, and STING/TBK1/IRF3 activation. HeLa cells stably expressing E.V. or TENT5A-3×Flag were stimulated with increasing doses of CT-DNA. (D) Western blot analysis of STING ubiquitination in HEK293T cells transfected with STING-Flag, Myc-ubiquitin, and increasing doses of TENT5A-3×HA plasmids. (E) Western blot analysis of STING ubiquitination in HEK293T cells co-transfected with TENT5A-3×HA, STING-Flag, and WT or mutant ubiquitin plasmids (Myc-Ub, Myc-Ub-K6, Myc-Ub-K11, Myc-Ub-K27, Myc-Ub-K29, Myc-Ub-K33, Myc-Ub-K48, Myc-Ub-K63). (F) Co-IP and Western blot analysis of the STING-TBK1 interaction in HEK293T cells transfected with STING-Flag, TBK1-3×Myc, and increasing doses of TENT5A-3×HA plasmids. (G) Western blot analysis of IRF3 subcellular localization in cytoplasmic and nuclear fractions from HeLa cells stably expressing E.V. or TENT5A-3×Flag, following HSV-1 (MOI = 1) infection or CT-DNA (1 μg/mL, 9 h) stimulation. Lamin B1 and β-tubulin served as nuclear and cytoplasmic markers, respectively. (H) Immunofluorescence imaging of IRF3 nuclear translocation in HeLa cells stably expressing TENT5A-3×Flag or E.V. after being stimulated with CT-DNA (1 μg/mL, 9 h). The percentage of cells with nuclear IRF3 was quantified in 50 cells per group. Scale bar, 10 μm. Data are presented as mean ± S.D. Two-tailed unpaired Student’s t-test; \*\*\*\**P <* 0.0001.

We next examined whether TENT5A affects STING oligomerization, a critical step for signal platform assembly. Semi-denaturing detergent agarose gel electrophoresis (SDD-AGE) analysis of HEK293T cells co-transfected with STING-Flag and increasing doses of TENT5A-3×HA demonstrated that TENT5A suppressed the formation of high-molecular-weight STING oligomers in a dose-dependent manner (Supplementary Figure 8A). This inhibitory effect was also observed when TBK1 was co-expressed, indicating that TENT5A acts upstream of TBK1-mediated amplification of STING oligomerization (Supplementary Figure 8B). To validate these findings at the endogenous level, we transfected HeLa cells with increasing doses of TENT5A-3×HA and performed SDD-AGE to assess endogenous STING oligomerization. Consistent with the overexpression system, endogenous STING oligomer formation was progressively reduced as TENT5A expression increased (Figure 3B). To further dissect whether TENT5A selectively inhibits higher-order oligomerization without disrupting the basal STING dimer, we used HeLa cells stably expressing TENT5A or empty vector (E.V.) and stimulated them with increasing doses of CT-DNA. Native-PAGE analysis revealed that TENT5A overexpression significantly reduced STING oligomer band intensity but had no discernible effect on STING dimer levels, establishing that TENT5A specifically inhibits higher-order oligomerization while leaving the basal dimeric state intact. In parallel, SDS-PAGE analysis confirmed that TENT5A overexpression markedly reduced phosphorylation of TBK1, IRF3, and STING without altering total protein levels (Figure 3C), consistent with the signaling data presented above.

Ubiquitination is an important post-translational modification that regulates STING activation, with K27-, K48-, and K63-linked polyubiquitin chains playing key roles in promoting STING signaling. To investigate whether TENT5A modulates STING ubiquitination, we co-transfected HEK293T cells with STING-Flag, Myc-tagged ubiquitin (Myc-Ub), and increasing doses of TENT5A-3×HA, and assessed total STING ubiquitination by immunoprecipitating with anti-Flag beads. TENT5A overexpression reduced total STING ubiquitination in a dose-dependent manner (Figure 3D). To identify the specific ubiquitin linkage types affected by TENT5A, we employed a panel of seven single-lysine ubiquitin mutants (Myc-Ub-K6, Myc-Ub-K11, Myc-Ub-K27, Myc-Ub-K29, Myc-Ub-K33, Myc-Ub-K48, and Myc-Ub-K63) as well as two linkage-specific mutants (Myc-Ub-K48R and Myc-Ub-K63R). Co-IP analysis showed that TENT5A selectively suppressed K48- and K63-linked ubiquitination of STING, with a particularly pronounced effect on K63-linked chains (Figure 3E, Supplementary Figure 8C).

A critical consequence of STING oligomerization is the recruitment and activation of TBK1. We therefore investigated whether TENT5A affects the STING-TBK1 interaction. Co-IP experiments in HEK293T cells co-transfected with STING-Flag, TBK1-3×Myc, and increasing doses of TENT5A-3×HA showed that TENT5A overexpression markedly disrupted the STING-TBK1 interaction in a dose-dependent manner (Figure 3F). This result is consistent with the observed reduction in TBK1 autophosphorylation and subsequent IRF3 phosphorylation, placing the inhibitory effect of TENT5A upstream of TBK1 recruitment.

IRF3, the central transcription factor of the cGAS-STING pathway, must undergo phosphorylation, dimerization, and nuclear translocation to drive IFN-I gene expression. Given that TENT5A inhibits STING oligomerization, ubiquitination, TBK1 recruitment, and IRF3 phosphorylation, we reasoned that it would also block IRF3 nuclear translocation as a downstream consequence. To test this, we performed subcellular fractionation in HeLa cells stably expressing TENT5A-3×Flag or E.V., followed by HSV-1 infection or CT-DNA stimulation. Western blot analysis of cytoplasmic and nuclear fractions showed that, under both stimulation conditions, IRF3 was substantially retained in the cytoplasmic fraction and correspondingly reduced in the nuclear fraction in TENT5A-overexpressing cells compared to E.V. controls; the purity of subcellular fractions was confirmed by the cytoplasmic marker β-tubulin and the nuclear marker Lamin B1 (Figure 3G). To corroborate these findings at the single-cell level, we performed immunofluorescence analysis in HeLa cells stably expressing TENT5A-3×Flag or E.V. following CT-DNA stimulation. In E.V. control cells, IRF3 underwent robust nuclear translocation upon CT-DNA stimulation, as evidenced by strong nuclear fluorescence signals. In contrast, IRF3 fluorescence remained predominantly cytoplasmic in TENT5A-overexpressing cells, with markedly reduced nuclear accumulation (Figure 3H). Quantification confirmed that the percentage of cells exhibiting nuclear IRF3 was significantly lower in the TENT5A group. Taken together, these results demonstrate that TENT5A suppresses IRF3 nuclear translocation as a downstream consequence of its inhibition of the STING–TBK1 signaling axis, and that TENT5A acts at multiple sequential steps of the STING activation cascade to achieve broad and robust suppression of IFN-I induction.

### TENT5A inhibits STING ER-Golgi translocation

Translocation of STING from the endoplasmic reticulum (ER) to the Golgi apparatus is a critical and obligatory step in cGAS-STING pathway activation^[6, 16]^. Having established that TENT5A resides at the ER at steady state and interacts with STING, we investigated whether TENT5A influences this key trafficking event. To first confirm the subcellular localization of exogenous TENT5A relative to STING during infection, we performed immunofluorescence analysis in HeLa cells expressing TENT5A-3×Flag following HSV-1 infection. TENT5A-3×Flag co-localized extensively with the ER marker Calnexin, whereas STING redistributed away from the ER upon infection; importantly, TENT5A remained anchored at the ER and did not redistribute with STING (Supplementary Figure 9A). Similarly, following Serum Starvation, TENT5A-3×HA maintained strong ER localization, while activated STING accumulated at peri-Golgi regions, indicating that the TENT5A–STING interaction is spatially confined to the ER (Supplementary Figure 9B). To corroborate these findings with endogenous proteins, we assessed the co-localization of endogenous TENT5A with Calnexin and endogenous STING in HeLa cells following HSV-1 infection. Endogenous TENT5A co-localized with Calnexin throughout the time course of infection, and its co-localization with STING progressively decreased as STING translocated away from the ER upon viral stimulation (Figure 4A). Together, these observations establish that TENT5A is an ER-resident protein that does not follow STING during its activation-induced trafficking.

**Figure 4.**
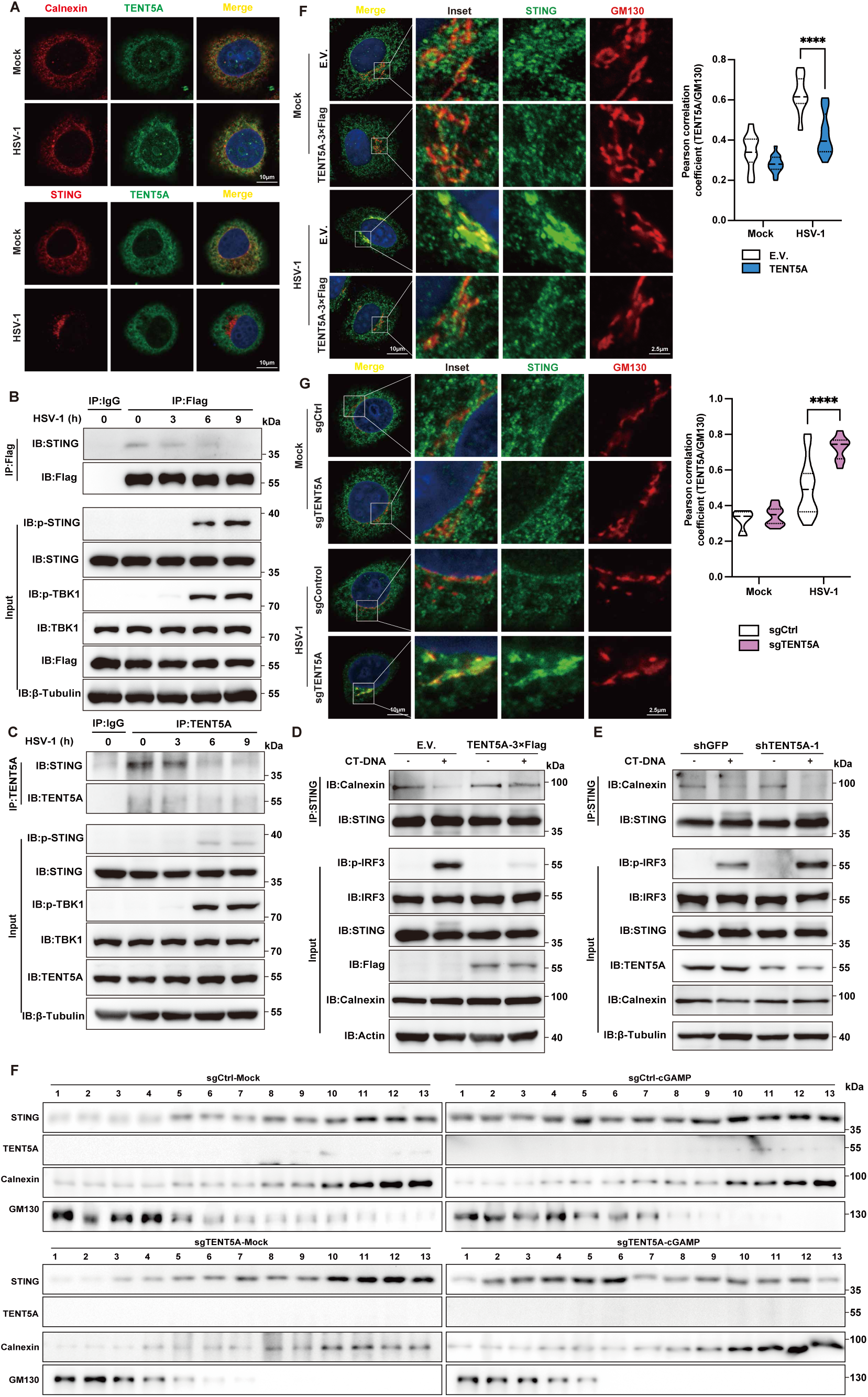
TENT5A inhibits STING translocation. (A) Immunofluorescence colocalization analysis of endogenous TENT5A with Calnexin (top)/STING (bottom) in HeLa cells after HSV-1 (MOI = 1) infection. Scale bar, 10 μm. (B, C) Co-IP and Western blot analysis of the STING-TENT5A interaction in TENT5A-expressing (B) or WT (C) HeLa cells at the indicated time points post HSV-1 (MOI = 1) infection. (D, E) Co-IP and Western blot analysis of the STING-Calnexin interaction in E.V./TENT5A-expressing THP-1 cells (D) or shGFP/shTENT5A THP-1 cells (E) following CT-DNA (1 μg/mL) stimulation. (F, G) HeLa cells stably expressing E.V./TENT5A-3×Flag (F) or sgCtrl/sgTENT5A (G) were mock-infected or infected with HSV-1 (MOI = 1, 5 h). Immunofluorescence staining was performed to assess STING colocalization with the Golgi marker GM130. Colocalization was quantified by Pearson’s correlation coefficient in 20 cells using ImageJ software. Scale bar, 10 μm (main image), 2.5 μm (inset). Data are presented as mean ± S.D. Two-tailed unpaired Student’s t-test; \*\*\*\**P <* 0.0001. (H) sgCtrl and sgTENT5A HeLa cells were stimulated with cGAMP (1 μg/mL, 2 h). Thirteen subcellular fractions (No. 1 to No. 13) were isolated via sucrose density gradient centrifugation. The endoplasmic reticulum marker Calnexin was gradually enriched, whereas the Golgi marker GM130 was gradually depleted from fraction 1 to fraction 13. The distribution of STING in the ER and Golgi compartments was detected.

The progressive spatial separation of TENT5A from activated STING prompted us to examine the dynamics of their physical interaction over the course of viral infection. Co-IP experiments in HeLa cells stably expressing TENT5A-3×Flag showed that the association between TENT5A and STING was robust at early time points after HSV-1 infection but diminished progressively as infection proceeded, correlating with the onset of STING translocation (Figure 4B). To confirm this at the endogenous level, we performed Co-IP using an anti-TENT5A antibody in wild-type HeLa cells at sequential time points post HSV-1 infection. Consistent with the exogenous data, the interaction between endogenous TENT5A and endogenous STING was strong at early time points and gradually weakened to near-undetectable levels as infection progressed (Figure 4C). These dynamic interaction data are fully consistent with a model in which TENT5A engages STING at the ER before its activation-induced departure, and the two proteins dissociate as STING exits the ER.

Based on these localization and interaction data, we hypothesized that TENT5A, by anchoring STING at the ER, impedes STING’s activation-induced translocation to the Golgi. To test this, we first examined the effect of TENT5A overexpression on STING’s association with ER and Golgi markers by immunofluorescence in HeLa cells stably expressing TENT5A-3×Flag or empty vector (E.V.) following HSV-1 infection. TENT5A overexpression significantly increased the co-localization of endogenous STING with the ER marker Calnexin compared to E.V. controls, indicating that TENT5A promotes ER retention of STING (Supplementary Figure 9C). Correspondingly, the co-localization of STING with the Golgi marker GM130 was markedly reduced in TENT5A-overexpressing cells, demonstrating that TENT5A blocks STING trafficking to the Golgi (Figure 4F). To validate the ER retention mechanism at the protein interaction level, we performed Co-IP experiments in E.V. and TENT5A-overexpressing THP-1 cells stimulated with CT-DNA. Immunoprecipitation of STING revealed a significantly stronger association between STING and Calnexin in TENT5A-overexpressing cells than in E.V. controls (Figure 4D), providing biochemical evidence that TENT5A enhances the anchorage of STING to the ER membrane and thereby prevents its translocation.

To complement the overexpression experiments with loss-of-function evidence, we assessed STING translocation in TENT5A-deficient cells. Immunofluorescence analysis of sgTENT5A knockout HeLa cells following HSV-1 infection revealed a significant decrease in STING–Calnexin co-localization compared to sgCtrl controls, indicating that loss of TENT5A facilitates STING departure from the ER (Supplementary Figure 9D). Concordantly, STING–GM130 co-localization was significantly elevated in sgTENT5A cells, demonstrating enhanced STING translocation to the Golgi upon TENT5A depletion (Figure 4G). To extend this finding to a physiologically relevant primary cell model, we examined STING translocation in *Tent5a*^fl/fl^ and *Tent5a*^fl/fl^ Lyz2-Cre BMDMs following HSV-1 infection. Immunofluorescence quantification showed that STING–Calnexin co-localization was markedly reduced in TENT5A-deficient BMDMs compared to controls (Supplementary Figure 9E), confirming that TENT5A restricts STING–ER dissociation in primary macrophages. At the biochemical level, Co-IP analysis of shTENT5A-knockdown THP-1 cells stimulated with CT-DNA showed that the STING–Calnexin interaction was substantially weakened upon TENT5A depletion (Figure 4E), consistent with the immunofluorescence findings and supporting the conclusion that TENT5A maintains STING–ER anchorage.

To further confirm TENT5A governs STING subcellular distribution via organelle separation, we performed sucrose density gradient ultracentrifugation to fractionate ER and Golgi compartments from sgCtrl and sgTENT5A HeLa cells treated with cGAMP (Figure 4H). Continuous discontinuous sucrose gradients (29%, 35%, 37% w/w) were subjected to ultracentrifugation, and 200 μL sequential fractions (1–13) were collected from the tube bottom. Proteins in each fraction were concentrated using methanol-chloroform precipitation and analyzed by immunoblotting. Following cGAMP stimulation, STING abundance in Golgi-enriched fractions increased markedly; this Golgi accumulation was further amplified in TENT5A-knockout cells, providing definitive biochemical proof that TENT5A depletion promotes STING trafficking from the ER to the Golgi apparatus.

Collectively, the gain-of-function and loss-of-function data from multiple cell types and experimental approaches converge to demonstrate that TENT5A inhibits the ER-to-Golgi translocation of STING by retaining it at the ER, thereby suppressing a pivotal licensing step required for full cGAS-STING pathway activation.

### TENT5A competitively inhibits the interaction between STING and SEC24B

Having established that TENT5A inhibits STING ER-to-Golgi translocation, we next sought to define the precise molecular mechanism by which it does so. STING export from the ER is mediated by COPII-coated vesicles, whose assembly requires the small GTPase SAR1 and the inner coat adaptors SEC23 and SEC24, together with the outer coat proteins SEC13 and SEC31^[17–19]^. To directly test whether TENT5A impairs STING packaging into COPII vesicles, we employed an in vitro COPII vesicle reconstitution assay (Figure 5A). Semi-permeabilized cells and cytosolic fractions prepared from sgCtrl or sgTENT5A HeLa cells pre-stimulated with cGAMP were incubated together to reconstitute cargo loading into nascent vesicles, using TMP21 as a COPII vesicle marker. STING packaging efficiency into COPII vesicles was significantly elevated in TENT5A-deficient cells compared to sgCtrl controls (Figure 5B), demonstrating that TENT5A suppresses the physical incorporation of STING into COPII vesicles and thereby restricts its ER export at a mechanistic level.

**Figure 5.**
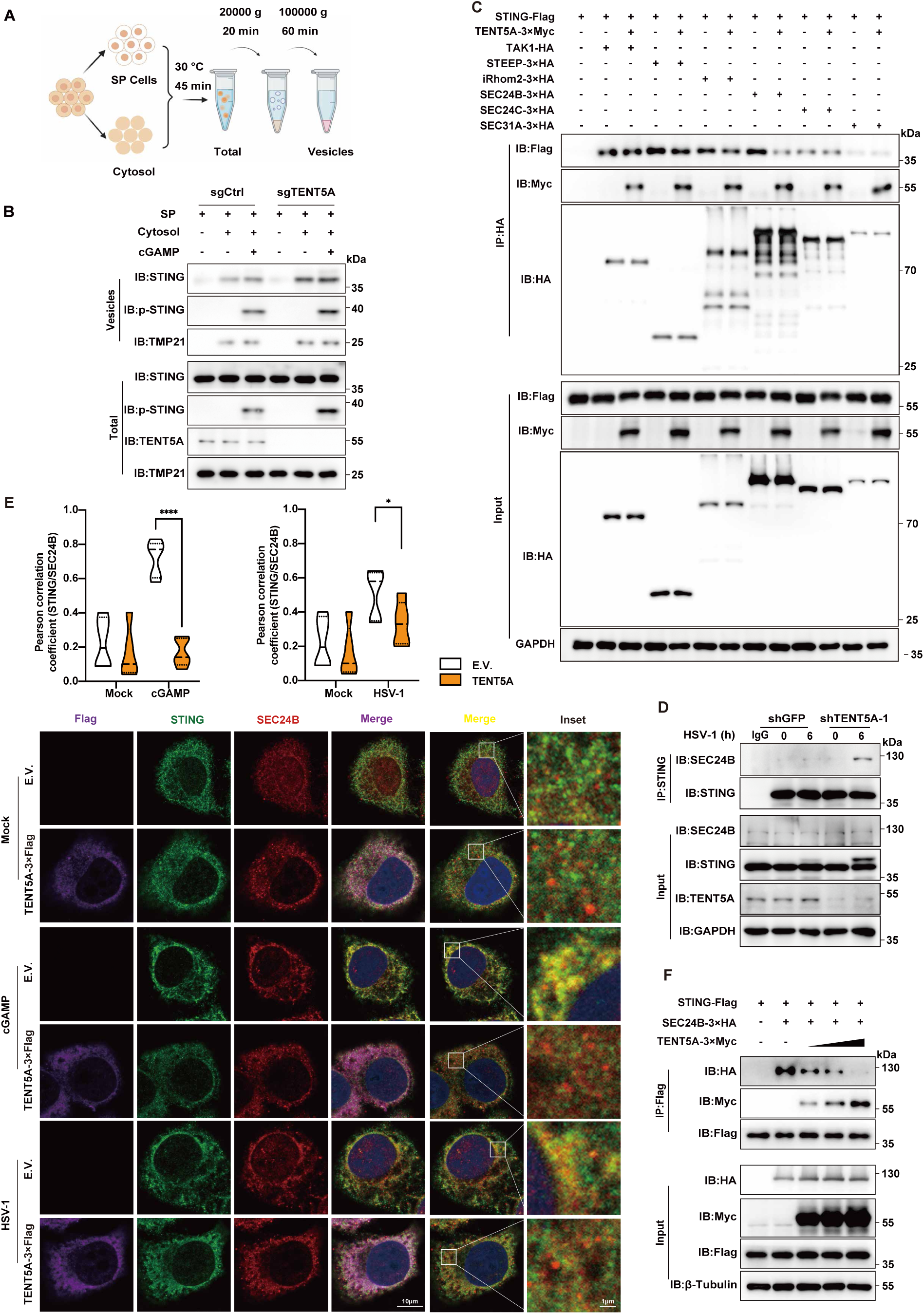
TENT5A competitively inhibits the interaction between STING and SEC24B. (A) Schematic diagram of the in vitro COPII vesicle reconstitution assay. (B) Western blot analysis of STING packaging into COPII vesicles in HeLa cells stably expressing sgCtrl or sgTENT5A, stimulated with cGAMP (1μg/mL, 2 h). TMP21 served as a COPII vesicle marker. (C) Co-IP was performed using anti-HA agarose beads with lysates from HEK293T cells expressing STING-HA, TENT5A-3×Myc, and plasmids encoding candidate STING trafficking-related proteins (TAK1-HA, STEEP-3×HA, iRhom2-3×HA, SEC24B-3×HA, SEC24C-3×HA, SEC31A-3×HA). (D) Co-IP was performed using anti-STING antibody with lysates from HeLa cells stably transfected with shGFP or shTENT5A and infected with HSV-1 (MOI = 1, 6 h). (E) Immunofluorescence analysis of endogenous STING-SEC24B colocalization. HeLa cells stably expressing E.V. or TENT5A were stimulated with cGAMP (1μg/mL, 0.5 h) or infected with HSV-1 (MOI = 1, 3 h). Colocalization was quantified by Pearson’s correlation coefficient in 20 cells using ImageJ software. Scale bars: 10 μm (main image), 1 μm (inset). Data are presented as mean ± S.D. Two-tailed unpaired Student’s t-test; \**P <* 0.05, \*\*\*\**P <* 0.0001. (F) Co-IP was performed using anti-Flag magnetic beads with lysates from HEK293T cells expressing STING-Flag, SEC24B-3×HA, and increasing amounts of TENT5A-3×Myc.

To identify the molecular intermediary through which TENT5A disrupts STING’s COPII-mediated ER export, we performed a systematic Co-IP screen examining two classes of proteins for their capacity to interact with both STING and TENT5A: known STING translocation regulators (TAK1, STEEP, iRhom2, TRAPβ, SEC61B, SEC5) and the full complement of COPII subunits (SAR1A, SAR1B, SEC23A, SEC23B, SEC24A, SEC24B, SEC24C, SEC24D, SEC13, SEC31A, SEC31B). Each candidate was assessed for interaction with STING-Flag (Supplementary Figure 10A, 10B) and with TENT5A-3×Myc (Supplementary Figure 10C, 10D) in HEK293T cells by Co-IP using anti-HA beads. Cross-referencing both datasets identified six proteins including TAK1, STEEP, iRhom2, SEC24B, SEC24C, and SEC31A, that interact with both STING and TENT5A, nominating them as candidates for a regulatory nexus. To determine which of these shared interactors is functionally disrupted by TENT5A, we performed a focused Co-IP experiment in HEK293T cells co-expressing STING-HA, TENT5A-3×Myc, and each of the six candidate proteins. TENT5A overexpression selectively and substantially reduced the interaction of STING with SEC24B, while leaving STING associations with TAK1, STEEP, iRhom2, SEC24C, and SEC31A largely intact (Figure 5C), pinpointing SEC24B as the critical node through which TENT5A disrupts STING’s COPII-mediated ER export.

To validate the regulation of the STING–SEC24B interaction by TENT5A through complementary loss-of-function approaches, we assessed STING–SEC24B binding in multiple TENT5A-knockdown models. First, in HEK293T cells stably expressing shGFP, shTENT5A-1, or shTENT5A-2, co-transfection of STING-Flag and SEC24B-3×HA followed by Co-IP with anti-HA beads showed that STING–SEC24B interaction was significantly enhanced in both knockdown lines compared to shGFP controls, confirming the specificity of the regulatory effect with two independent shRNAs (Supplementary Figure 11A). Extending this to endogenous proteins, Co-IP with an anti-STING antibody in shTENT5A-1 HeLa cells stimulated with CT-DNA demonstrated a marked increase in endogenous STING–SEC24B association compared to shGFP controls (Supplementary Figure 11B). To further examine the temporal dynamics of this regulation, we performed time-course Co-IP experiments in shGFP and shTENT5A-1 THP-1 cells stimulated with cGAMP. The STING–SEC24B interaction was progressively enhanced in shTENT5A-1 cells relative to controls across all time points examined, with the enhancement becoming more pronounced as stimulation time increased — a pattern consistent with the kinetics of STING translocation (Supplementary Figure 11C). Complementing these knockdown data, Co-IP analysis in shTENT5A HeLa cells following HSV-1 infection likewise showed significantly elevated endogenous STING–SEC24B binding upon TENT5A depletion (Figure 5D), collectively establishing across three cell types and multiple stimuli that endogenous TENT5A constitutively restrains STING–SEC24B complex formation during innate immune activation.

To corroborate these biochemical findings at the spatial level, we examined the co-localization of endogenous STING and SEC24B by immunofluorescence in HeLa cells stably expressing TENT5A-3×Flag or E.V., subjected to cGAMP stimulation or HSV-1 infection. Quantification by Pearson’s correlation coefficient showed that TENT5A overexpression markedly reduced the spatial overlap between STING and SEC24B under both stimulation conditions compared to E.V. controls, with the co-localization signal significantly diminished in TENT5A-expressing cells (Figure 5E). The spatial disruption of the STING–SEC24B interaction by TENT5A is fully consistent with the biochemical Co-IP data across the knockdown models and reinforces the model that TENT5A prevents STING from engaging the COPII sorting adaptor SEC24B at the ER membrane, thereby blocking its ER export.

To determine whether TENT5A inhibits STING–SEC24B complex formation through a direct competitive mechanism, we performed a dose-dependent competition Co-IP assay in HEK293T cells co-transfected with STING-Flag and SEC24B-3×HA in the presence of increasing amounts of TENT5A-3×Myc. Immunoprecipitation with anti-Flag beads revealed that the STING–SEC24B interaction was progressively diminished as TENT5A expression increased, demonstrating dose-dependent competitive inhibition (Figure 5F). These results indicate that TENT5A and SEC24B compete for binding to an overlapping or mutually exclusive site on STING, and that elevated TENT5A occupancy of STING sterically or allosterically precludes SEC24B recruitment. Taken together, the data in this section establish a mechanistic model in which TENT5A, engages the CTT of STING and competitively displaces the COPII cargo adaptor SEC24B, thereby blocking STING packaging into COPII vesicles, preventing its ER-to-Golgi translocation, and ultimately suppressing cGAS-STING-mediated IFN-I production.

### TENT5A dampens the host immune defense against DNA viruses

The mechanistic studies presented above demonstrate that TENT5A suppresses cGAS-STING-mediated IFN-I signaling by blocking STING ER-to-Golgi translocation, thereby facilitating intracellular DNA virus replication. Consistent with this, TENT5A overexpression augmented *HSV-1-TK* mRNA levels (Supplementary Figure 4C, while TENT5A knockdown or knockout reduced *HSV-1-TK* (Figure 1A, F:Supplementary Figure 2B, 5C, 6A) and *VACV-J2* (Supplementary Figure 3C) mRNA levels and diminished viral protein ICP0 (Figure 1D, H: Supplementary Figure 3E, 5E, 6C) expression across multiple cell-based models. To directly quantify the impact of TENT5A on viral replication, we performed plaque assays on culture supernatants harvested from HSV-1-infected cells. In HeLa cells stably expressing sgCtrl, sgTENT5A-1, or sgTENT5A-2, TENT5A knockout resulted in a significant reduction in HSV-1 viral titers at 24 h post-infection compared to sgCtrl controls, with consistent results across both independent sgRNA lines (Supplementary Figure 12A). Likewise, in BMDMs derived from *Tent5a*^fl/fl^ and *Tent5a*^fl/fl^ Lyz2-Cre mice, TENT5A deficiency significantly reduced HSV-1 titers in cell culture supernatants following infection (Supplementary Figure 12B). These results establish that TENT5A promotes intracellular DNA virus replication in both human and murine cell types, and that its loss substantially enhances the antiviral capacity of cells.

To determine whether TENT5A exerts a physiologically relevant regulatory role in antiviral innate immunity in vivo, we established an HSV-1 infection model in littermate *Tent5a*^fl/fl^ and *Tent5a*^fl/fl^ Lyz2-Cre mice. Following intravenous administration of HSV-1, tissues and blood were collected at 72 h post-infection for assessment of antiviral gene expression. qPCR analysis of liver, spleen, lung, brainstem, and peripheral blood revealed that *Ifnb1* and *Cxcl10* mRNA levels were significantly elevated across all tissues examined in *Tent5a*^fl/fl^ Lyz2-Cre mice compared to *Tent5a*^fl/fl^ controls, while *HSV-1-TK* mRNA, a proxy for viral replication, was correspondingly reduced, indicating that myeloid-specific TENT5A deficiency enhances systemic antiviral IFN-I responses while suppressing viral propagation in vivo (Figure 6A). Corroborating these transcriptional findings, serum IFN-β protein levels measured by ELISA at 9 h post-infection were significantly higher in *Tent5a*^fl/fl^ Lyz2-Cre mice than in *Tent5a*^fl/fl^ littermates (Figure 6B), confirming that TENT5A restricts systemic IFN-β production during acute DNA virus infection in vivo.

**Figure 6.**
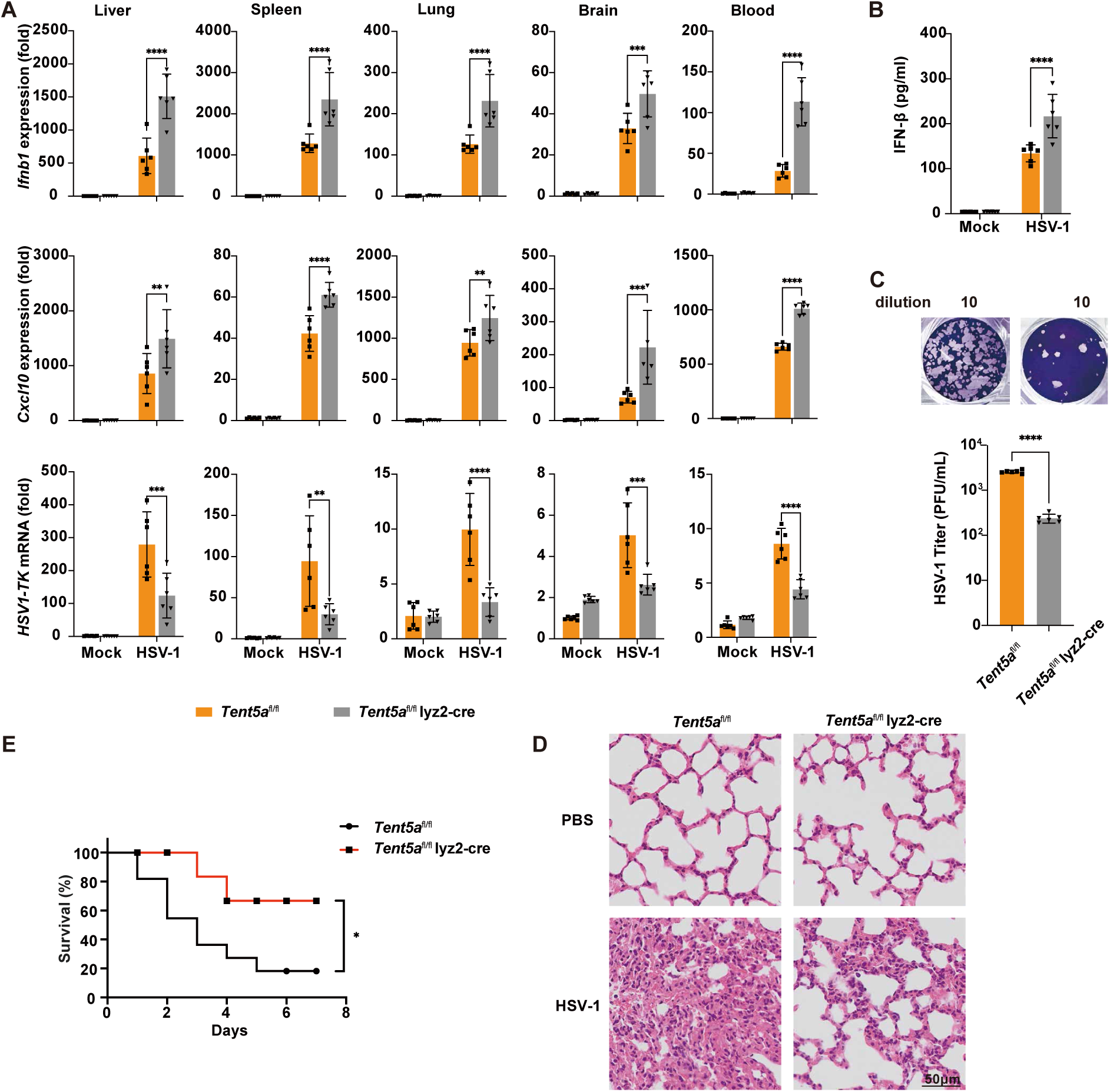
*Tent5a*-deficient Mice Exhibit Enhanced Tolerance to DNA Virus Infection. (A) Littermate *Tent5a*^fl/fl^ and *Tent5a*^fl/fl^ Lyz2-Cre mice (n = 6) were intravenously administered with HSV-1 (1×10^7^ PFU per mouse). The mRNA levels of *Ifnb1, Cxcl10*, and *HSV-1-TK* in liver, spleen, lung, brainstem, and peripheral blood were measured by qPCR at 72 h post-infection. (B) Littermate *Tent5a*^fl/fl^ and *Tent5a*^fl/fl^ Lyz2-Cre mice (n = 6) were intravenously administered with HSV-1 (1×10^7^ PFU per mouse) and serum IFN-β levels were detected by ELISA at 9 h post-infection. (C) Littermate *Tent5a*^fl/fl^ and *Tent5a*^fl/fl^ Lyz2-Cre mice (n = 6) were intravenously administered with HSV-1 (1×10^7^ PFU per mouse). Viral titers in liver tissues were determined by plaque assays at 72 h post-infection. (D) Littermate *Tent5a*^fl/fl^ and *Tent5a*^fl/fl^ Lyz2-Cre mice (n = 3) were intravenously administered with HSV-1 (1×10^7^ PFU per mouse, 72 h). Lung tissues were harvested for Hematoxylin and eosin (H&E) staining at 72 h post-infection. Scale bar, 50 μm. (E) Littermate *Tent5a*^fl/fl^ and *Tent5a*^fl/fl^ Lyz2-Cre mice (n = 10) were intravenously administered with HSV-1 (1×10^8^ PFU per mouse), and survival was monitored daily for 7 days. Data are presented as mean ± S.D. Two-tailed Student’s t-test (A, B, C); Log-rank test (E); \**P <* 0.05, \*\**P <* 0.01, \*\*\**P <* 0.001, \*\*\*\**P <* 0.0001 (A, B, E).

To assess the functional consequences of enhanced IFN-I signaling in TENT5A-deficient mice on viral control and disease outcome, we examined viral burden, tissue pathology, and survival. Plaque assays performed on liver tissue homogenates at 72 h post-infection revealed that viral titers were significantly lower in *Tent5a*^fl/fl^ Lyz2-Cre mice compared to *Tent5a*^fl/fl^ controls (Figure 6C), demonstrating that myeloid TENT5A deficiency confers enhanced in vivo viral clearance. Histopathological examination of lung sections by hematoxylin and eosin (H&E) staining at 72 h post-infection showed markedly reduced inflammatory infiltration and tissue damage in *Tent5a*^fl/fl^ Lyz2-Cre mice relative to *Tent5a*^fl/fl^ littermates (Figure 6D), indicating that stronger innate immune activation in the absence of TENT5A is associated with improved tissue integrity rather than exacerbated immunopathology. Finally, survival analysis following a lethal HSV-1 challenge (1×10^8^ PFU per mouse) demonstrated that *Tent5a*^fl/fl^ Lyz2-Cre mice exhibited significantly improved survival over a 7-day monitoring period compared to *Tent5a*^fl/fl^ controls (Figure 6E). Taken together, these in vivo data demonstrate that myeloid-specific TENT5A deficiency augments antiviral IFN-I production, reduces viral replication and tissue damage, and improves host survival during DNA virus infection, establishing TENT5A as a physiologically relevant negative regulator of antiviral innate immunity that impairs the host’s ability to combat DNA virus challenge through suppression of the cGAS-STING pathway.

### Tent5a deficiency inhibits tumorigenesis and progression in mice

Besides its critical role in antiviral immunity, the cGAS-STING axis serves as a core pathway governing antitumor immune surveillance, which drives production of type I interferons and proinflammatory cytokines to bridge innate and adaptive antitumor responses. Since TENT5A suppresses cGAS-STING signaling in myeloid cells, we explored whether loss of TENT5A could boost in vivo antitumor immunity.

We established a syngeneic B16-Luc melanoma subcutaneous tumor model. Littermate *Tent5a*^fl/fl^ and *Tent5a*^fl/fl^ Lyz2-Cre mice were injected with B16-Luc cells into the dorsal flank, and in vivo bioluminescence imaging was performed on day 8 and day 14 after tumor inoculation. Representative bioluminescence signals were markedly weaker in *Tent5a*^fl/fl^ Lyz2-Cre mice relative to *Tent5a*^fl/fl^ littermate controls (Figure 7A). Quantitative analysis of total bioluminescence flux further confirmed restrained tumor growth in myeloid-specific Tent5a-deficient mice throughout the observation window (Figure 7B), demonstrating that myeloid TENT5A ablation impedes melanoma initiation and progression in vivo.

**Figure 7.**
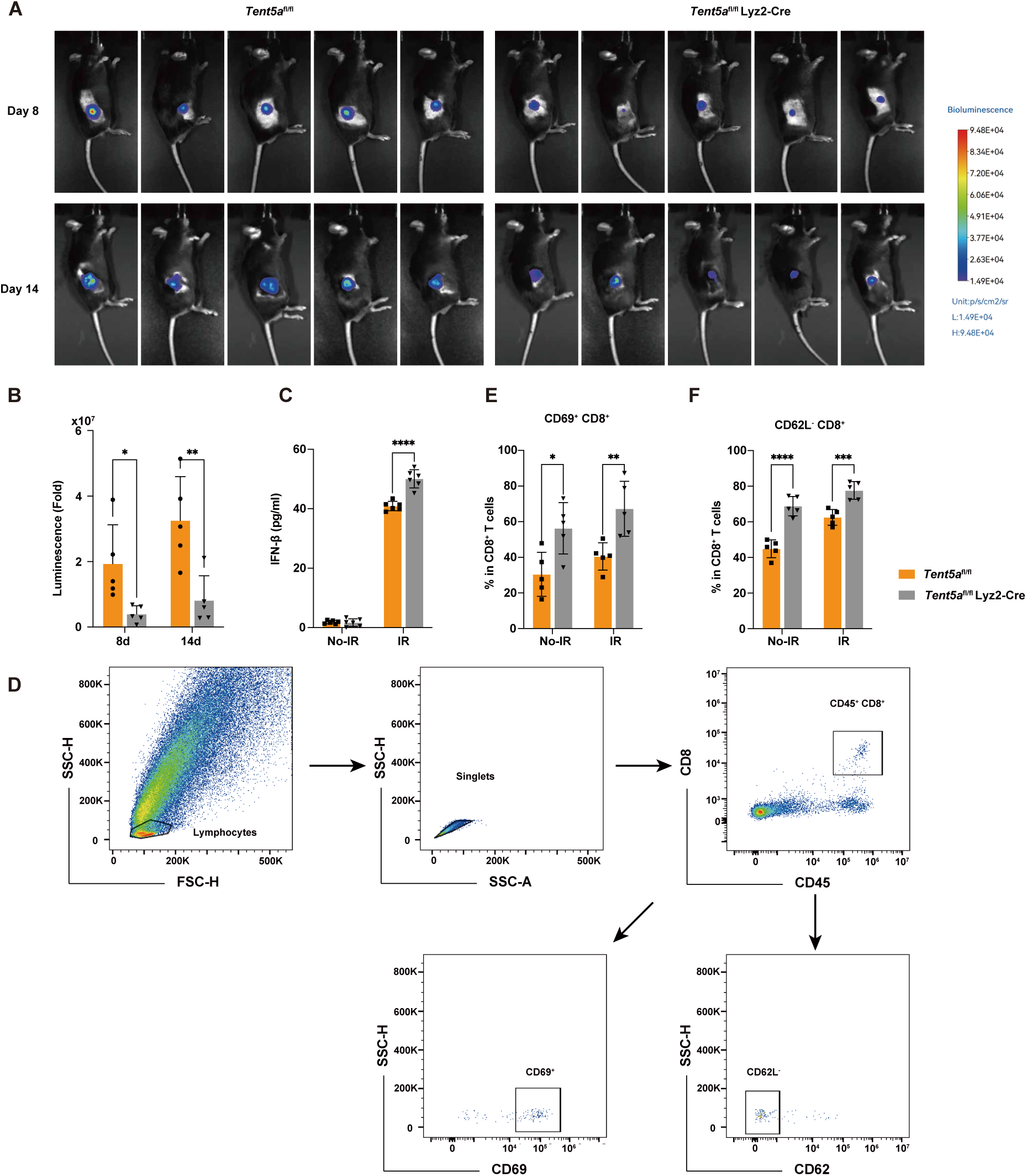
Tent5a-deficiency inhibits Tumorigenesis and Progression in mice. (A) Bioluminescent images of tumors were acquired in *Tent5a*^fl/fl^ and *Tent5a*^fl/fl^ Lyz2-Cre mice (n = 5) implanted with B16-Luc cells (5×10^5^ cells per mouse). (B) Quantification of tumor bioluminescence in mice as in (A) at the indicated time points. (C) Littermate *Tent5a*^fl/fl^ and *Tent5a*^fl/fl^ Lyz2-Cre mice (n = 6) were subcutaneously inoculated with B16-Luc cells (5×10^5^ cells per mouse) at the dorsal side. At 7 days post-inoculation, mice were irradiated at 800 cGy (current: 13.3 mA, dose rate: 300 cGy/min). 7 days post-irradiation serum IFN-β levels in mice were measured by ELISA. (D) Gating strategy for detecting the activation status of CD8⁺ T cells by flow cytometry. (E, F) Tumor tissues were harvested from mice as described in (D), homogenized, and processed into single-cell suspensions. The proportion of CD69^+^ CD8⁺ T cells (E) or CD62L^-^ CD8^+^T cells (F) were analyzed by flow cytometry. Data are presented as mean ± S.D. Two-tailed Student’s t-test; \**P <* 0.05, \*\**P <* 0.01, \*\*\**P <* 0.001, \*\*\*\**P <* 0.0001 (B, C, E, F).

**Figure 8.**
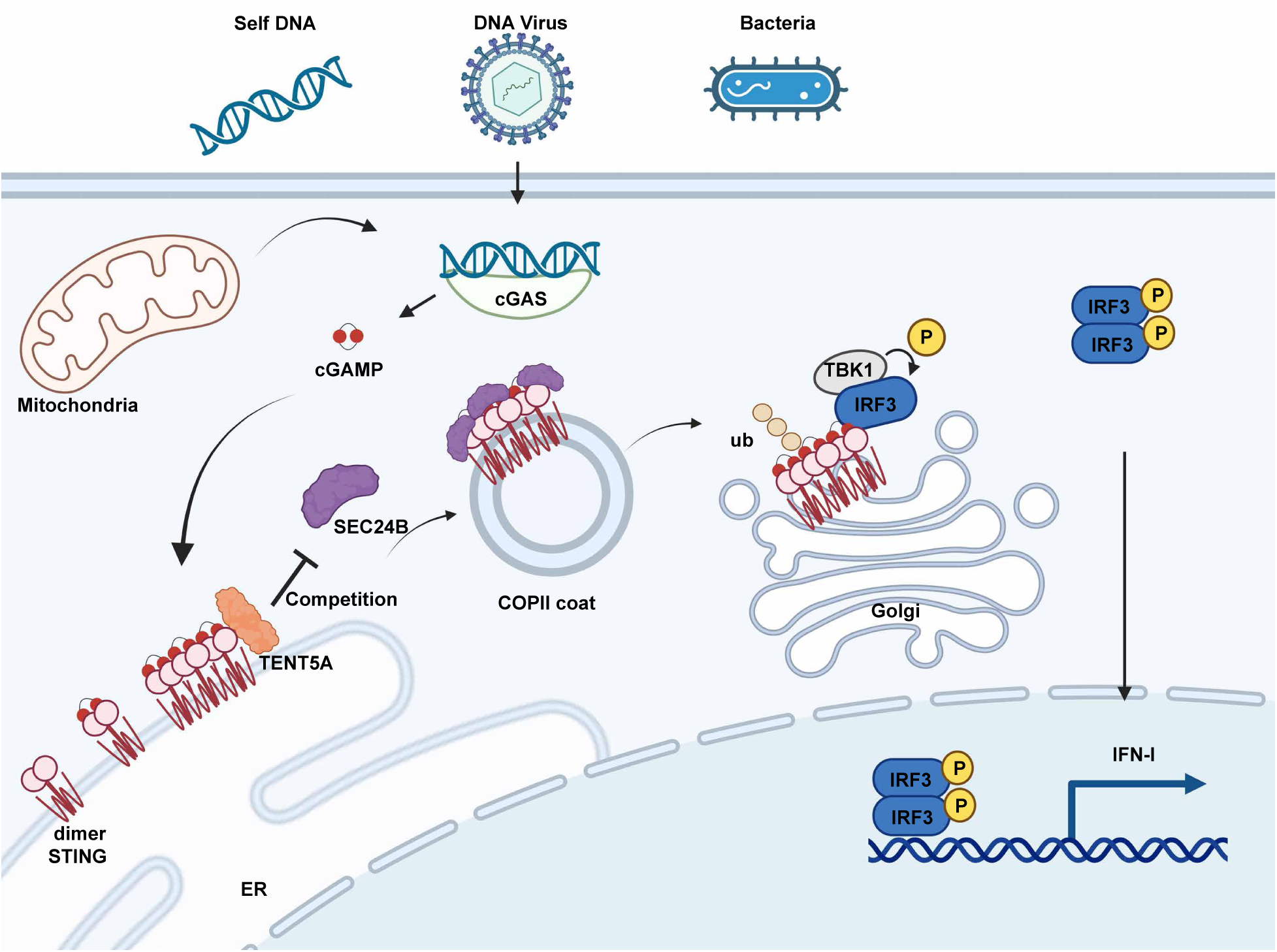
**A proposed model of TENT5A-mediated regulation of STING intracellular trafficking and innate immune activation.**

To further dissect the immunological mechanism by which TENT5A restrains tumor growth, we utilized an ionizing radiation model to amplify intratumoral cGAS-STING activation. Mice bearing subcutaneous B16-Luc tumors received whole-body irradiation (800 cGy, 13.3 mA, dose rate 300 cGy/min) on day 7 post inoculation. Seven days after radiation treatment, serum samples were collected for ELISA detection of IFN-β. The basal serum IFN-β level was low in non-irradiated tumor-bearing mice, while irradiation markedly induced IFN-β secretion; notably, *Tent5a*^fl/fl^ Lyz2-Cre mice exhibited significantly higher serum IFN-β concentrations than control mice under both irradiated and non-irradiated conditions (Figure 7C).

We next prepared single-cell suspensions from isolated tumor tissues and analyzed the activation and differentiation status of intratumoral CD8^+^ T cells via flow cytometry. The gating strategy for CD8^+^ T cell subsets was shown in Figure 7D. The frequency of CD69^+^activated CD8^+^ T cells was significantly elevated in tumors from *Tent5a*^fl/fl^ Lyz2-Cre mice (Figure 7E). Consistently, the proportion of CD62L^-^ effector CD8^+^ T cells was also obviously increased (Figure 7F). These data indicate that TENT5A deletion facilitates the differentiation of naive CD8^+^ T cells into activated effector subsets within the tumor microenvironment and reinforces tumor-specific cellular immunity.

Collectively, myeloid TENT5A loss potentiates cGAS-STING-dependent IFN-β production, promotes the activation and effector differentiation of intratumoral CD8^+^T cells, and robustly suppresses melanoma progression in vivo. Our results suggest that TENT5A represents a promising candidate target for tumor immunotherapy.

## Supplementary Figure

**Supplementary Figure 1.**
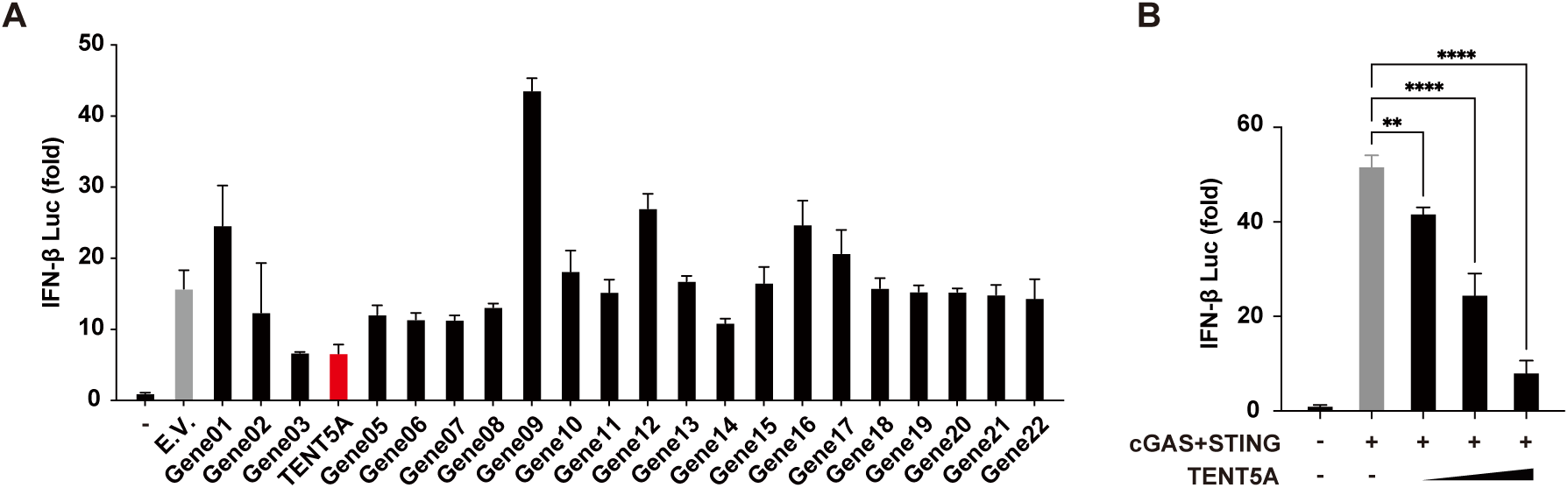
Dual-luciferase screening of candidate regulators of the cGAS-STING signaling pathway. (A) HEK293T cells stably expressing STING were co-transfected with expression constructs of cGAS, Gene01 to Gene22, and the IFN-β-Luc reporter plasmid. pRL-TK was used as the transfection internal control. Dual-luciferase assay were performed to quantify relative IFN-β promoter activity. (B) HEK293T cells stably expressing STING were co-transfected with expression constructs of cGAS, increasing amounts of TENT5A, and the IFN-β-Luc reporter plasmid. pRL-TK was used as the transfection internal control. Dual-luciferase assay were performed to verify the dose-dependent inhibitory effect of TENT5A on cGAS-STING signaling. Data are presented as mean ± S.D. Two-tailed unpaired Student’s t-test; \*\**P* < 0.01, \*\*\*\**P* < 0.0001. *P < 0.05.

**Supplementary Figure 2.**
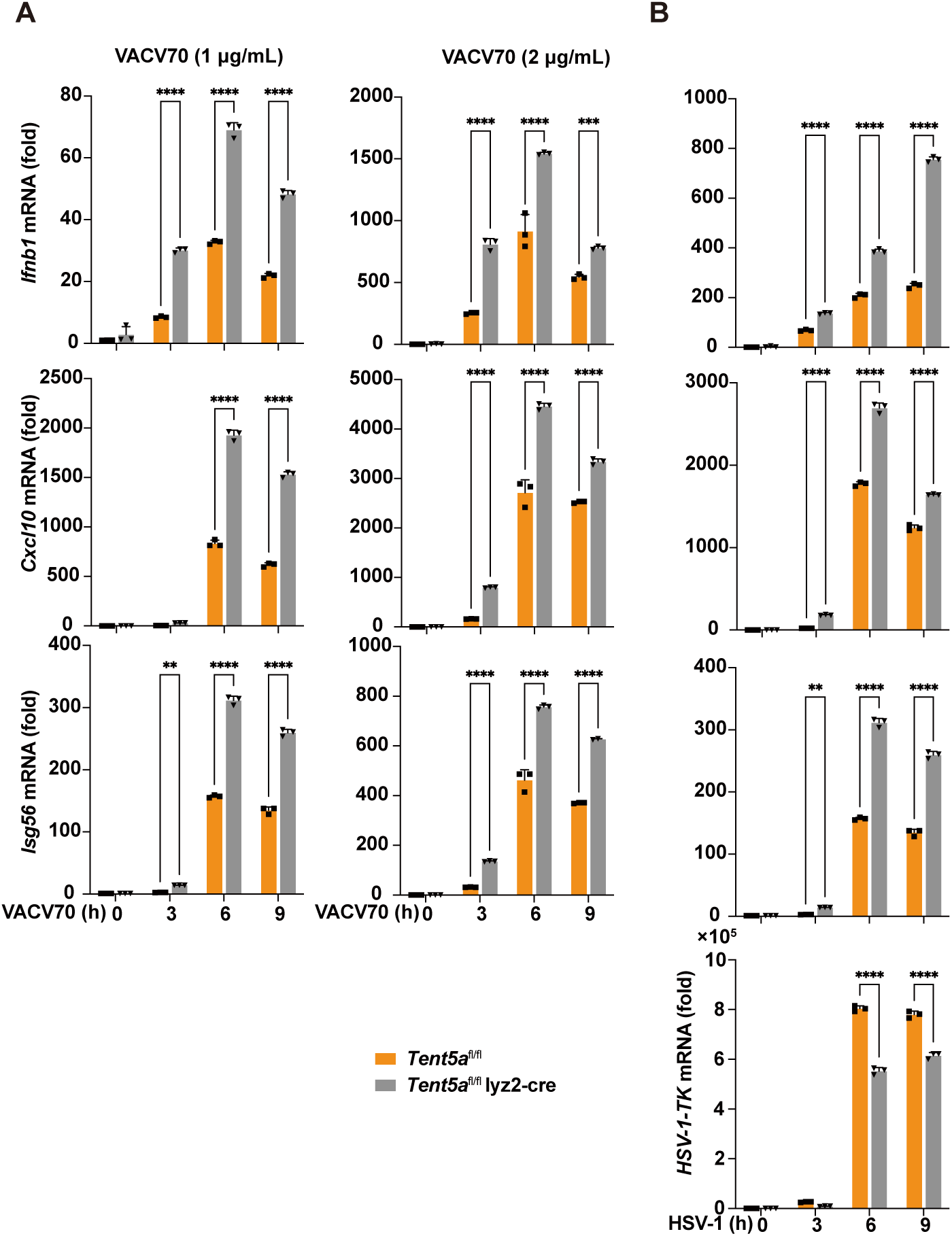
*Tent5a* deficiency promotes cGAS-STING-mediated IFN-I response in BMDMs. (A) qPCR analysis of *Ifnb1, Cxcl10,* and *Isg56* mRNA in *Tent5a*^fl/fl^ and *Tent5a*^fl/fl^ Lyz2-Cre BMDMs following stimulation with stimulation with VACV70 (1 μg/mL, 2 μg/mL) for the indicated time points. (B) qPCR analysis of *Ifnb1, Cxcl10, Isg56,* and *HSV-1-TK* mRNA in *Tent5a*^fl/fl^ and *Tent5a*^fl/fl^ Lyz2-Cre BMDMs following infection with HSV-1 (MOI = 2) for the indicated time points. Data are presented as mean ± S.D. Two-way ANOVA; \*\**P <* 0.01, \*\*\**P <* 0.001, \*\*\*\**P <* 0.0001 (A, B).

**Supplementary Figure 3.**
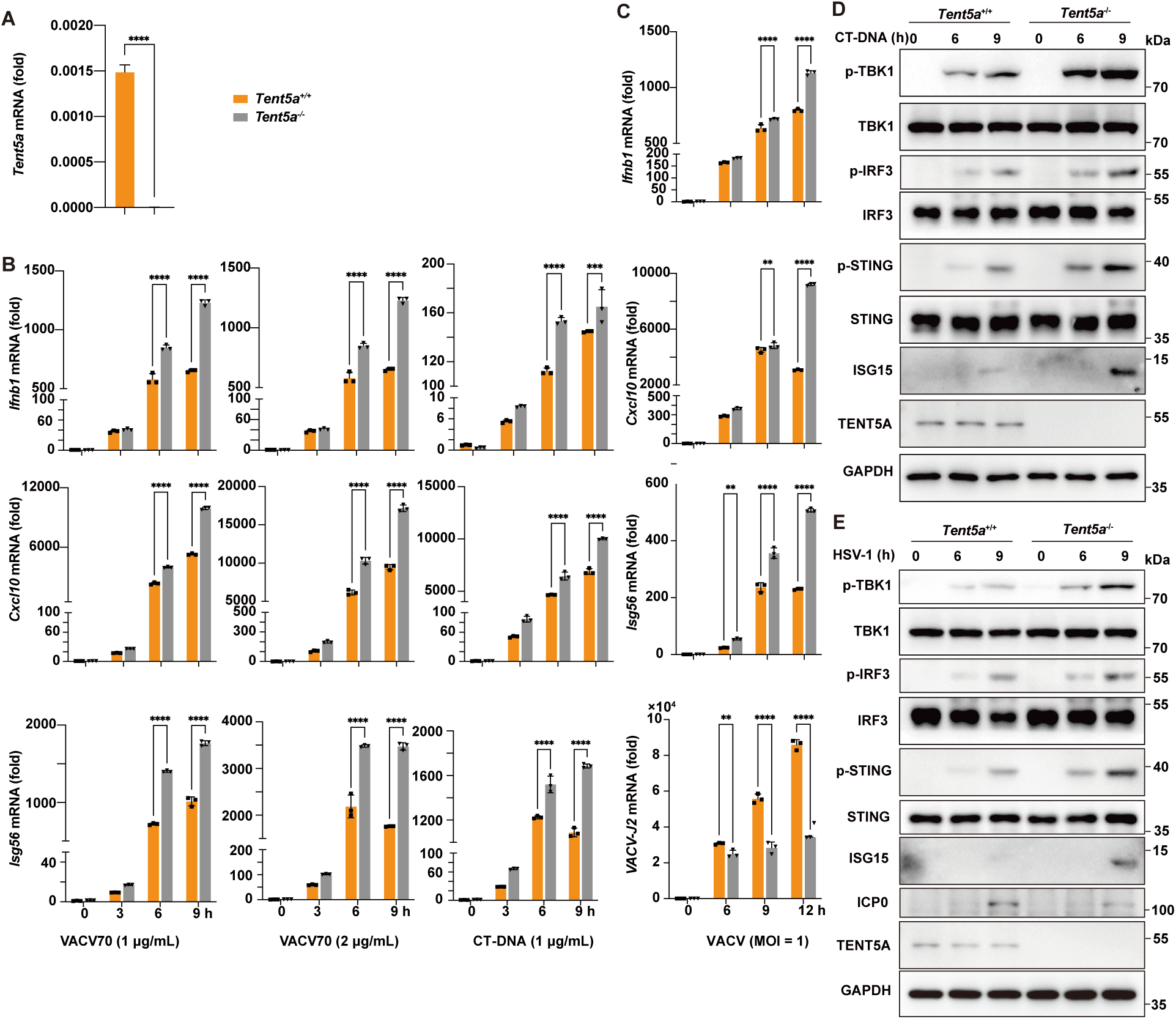
TENT5A deficiency promotes cGAS-STING-mediated IFN-I response in MEFs. (A) qPCR analysis of *Tent5a* mRNA in *Tent5a*^+/+^ and *Tent5a*^-/-^ MEFs. (B) qPCR analysis of *Ifnb1*, *Cxcl10*, and *Isg56* mRNA in *Tent5a*^+/+^ and *Tent5a*^-/-^ MEFs following stimulation with VACV70 (1 μg/mL, 2 μg/mL) or CT-DNA (1 μg/mL) for the indicated time points. (C) qPCR analysis of *Ifnb1*, *Cxcl10*, *Isg56*, and *VACV-J2* mRNA in *Tent5a*^+/+^ and *Tent5a*^-/-^ MEFs, following infection with VACV (MOI = 1) for the indicated time points. (D) Western blot analysis of ISG15 and phosphorylated and total STING, IRF3, and TBK1 in lysates of *Tent5a*^+/+^ and *Tent5a*^-/-^ MEFs stimulated with CT-DNA (1 μg/mL) for the indicated time points. (E) Western blot analysis of ISG15, ICP0, and phosphorylated and total STING, IRF3, and TBK1 in lysates of *Tent5a*^+/+^ and *Tent5a*^-/-^ MEFs infected with HSV-1 (MOI = 1) for the indicated time points. Data are presented as mean ± S.D. Two-way ANOVA; \*\**P <* 0.01, \*\*\**P <* 0.001, \*\*\*\**P <* 0.0001 (A, B, C).

**Supplementary Figure 4.**
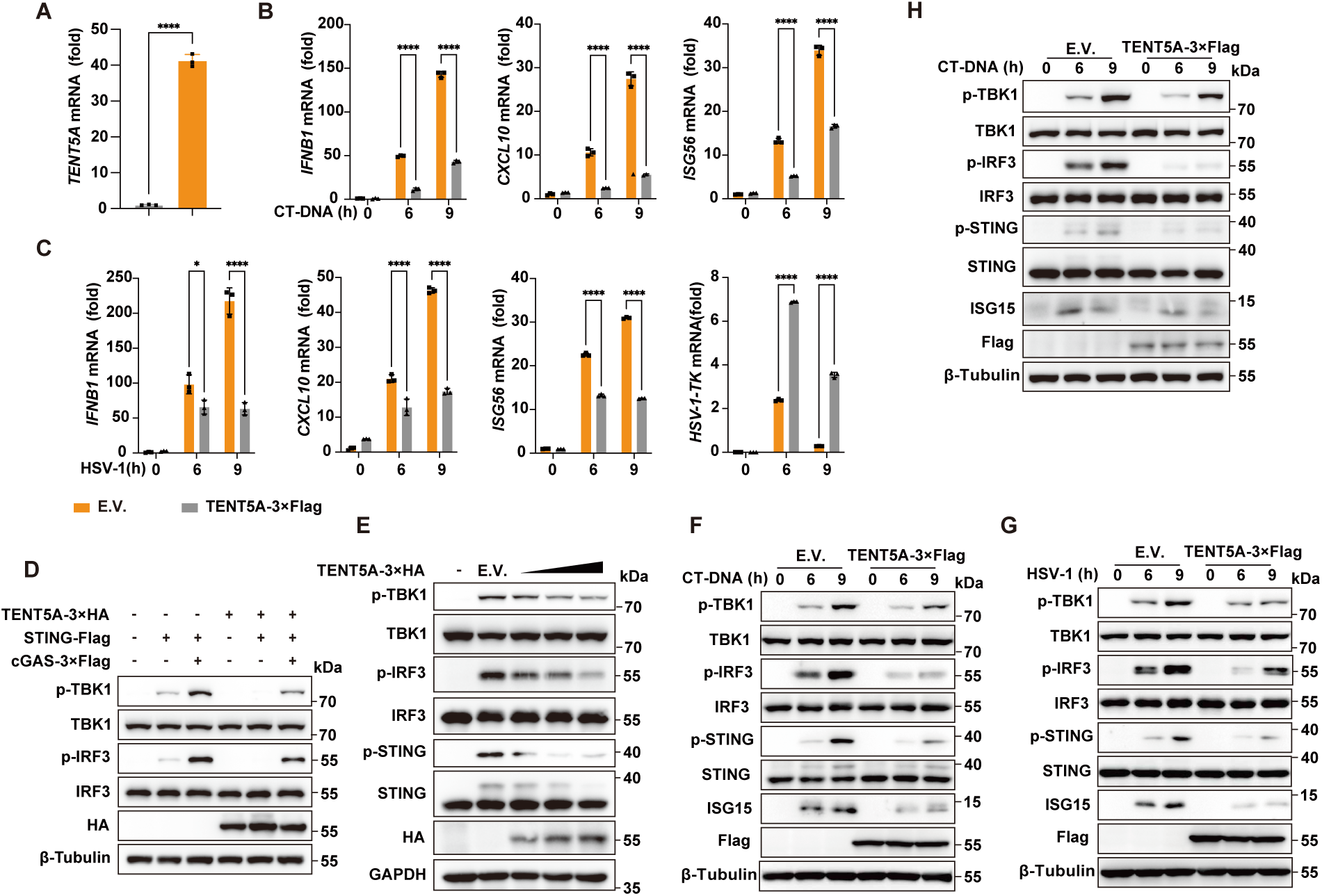
TENT5A-overexpression inhibits cGAS-STING-mediated IFN-I response. (A) qPCR analysis of *Tent5a* mRNA in HeLa cells stably expressing TENT5A-3×Flag or E.V. (B) qPCR analysis of *IFNB1*, *CXCL10*, and *ISG56* mRNA in HeLa cells stably expressing TENT5A-3×Flag or E.V. following stimulation with CT-DNA (1 μg/mL) for the indicated time points. (C) qPCR analysis of *IFNB1*, *CXCL10*, *ISG56*, and *HSV-1-TK* mRNA in HeLa cells stably expressing TENT5A-3×Flag or E.V. following infection with HSV-1 (MOI = 1) for the indicated time points. (D) Western blot analysis of phosphorylated and total IRF3 and TBK1 in HEK293T cells expressing TENT5A-3×HA, STING-Flag and cGAS-3×Flag. (E) Western blot analysis of phosphorylated and total STING, IRF3, and TBK1 in HeLa cells expressing gradient dose of TENT5A-3×HA. (F) Western blot analysis of ISG15 as well as phosphorylated and total STING, TBK1 and IRF3 in HeLa cells stably expressing TENT5A-3×Flag or E.V. following stimulated with CT-DNA (1 μg/mL) for the indicated time. (G) Western blot analysis of ISG15 as well as phosphorylated and total STING, TBK1 and IRF3 in HeLa cells stably expressing TENT5A-3×Flag or E.V. following infected with HSV-1 (MOI = 1) for the indicated time. (H) Western blot analysis of ISG15 as well as phosphorylated and total STING, TBK1 and IRF3 in THP-1 cells stably expressing TENT5A-3×Flag or E.V. following stimulated with CT-DNA (1 μg/mL) for the indicated time. Data are presented as mean ± S.D. Two-way ANOVA; \**P <* 0.05, \*\*\*\**P <* 0.0001 (A, B, C).

**Supplementary Figure 5.**
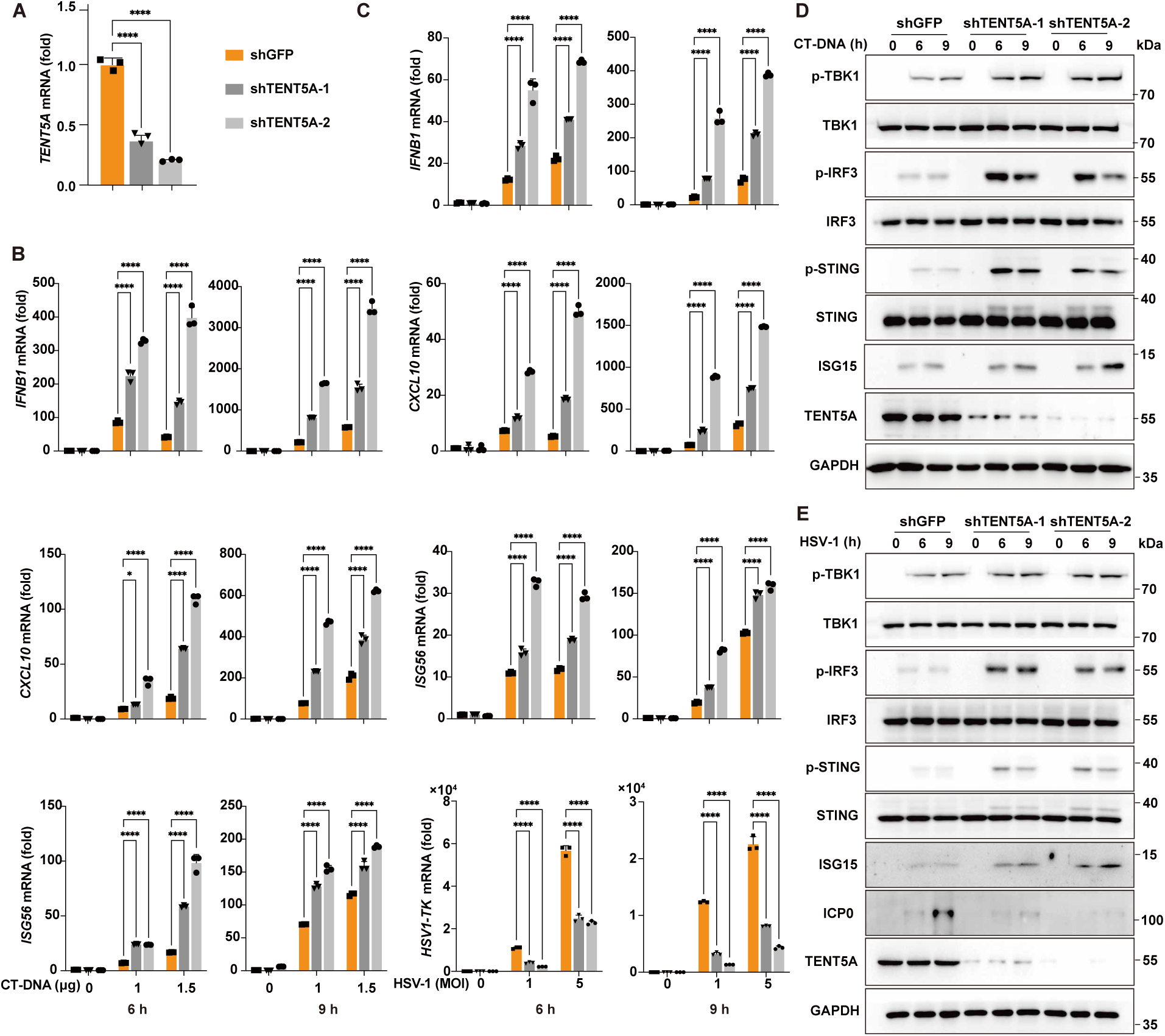
TENT5A knockdown promotes cGAS-STING-mediated IFN-I response. (A) qPCR analysis of *Tent5a* mRNA in THP-1 cells stably expressing shGFP, shTENT5A-1, and shTENT5A-2. (B) qPCR analysis of *IFNB1, CXCL10*, and *ISG56* mRNA in THP-1 cells stably expressing shGFP, shTENT5A-1, and shTENT5A-2 following stimulation with CT-DNA (1 μg/mL, 1.5 μg/mL) for the indicated time points. (C) qPCR analysis of *IFNB1, CXCL10, ISG56,* and *HSV-1-TK* mRNA in THP-1 cells stably expressing shGFP, shTENT5A-1, and shTENT5A-2 following infection with HSV-1 (MOI = 1, MOI = 5) for the indicated time points. (D) Western blot analysis of ISG15 as well as phosphorylated and total STING, TBK1 and IRF3 in THP-1 cells stably expressing shGFP, shTENT5A-1, and shTENT5A-2 following stimulation with CT-DNA (1 μg/mL) for the indicated time. (E) Western blot analysis of ISG15, ICP0, as well as phosphorylated and total STING, TBK1 and IRF3 in THP-1 cells stably expressing shGFP, shTENT5A-1, and shTENT5A-2 following infection with HSV-1 (MOI = 1) for the indicated time. Data are presented as mean ± S.D. Two-way ANOVA; \**P <* 0.05, \*\*\*\**P <* 0.0001 (A, B, C).

**Supplementary Figure 6.**
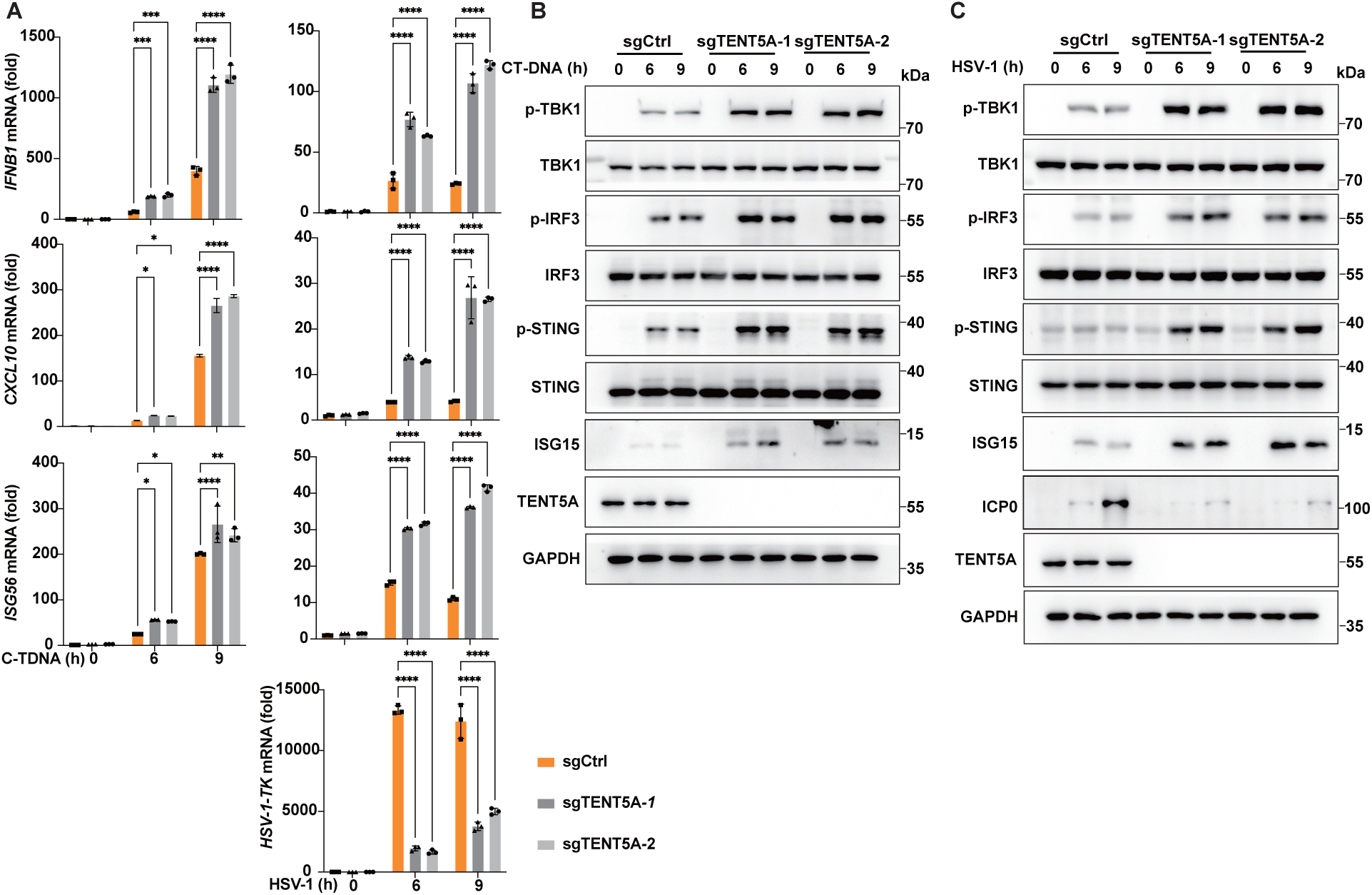
TENT5A knockout promotes cGAS-STING-mediated IFN-I response. (A) qPCR analysis of *IFNB1, CXCL10*, *ISG56*, and *HSV-1-TK* mRNA in THP-1 cells stably expressing sgCtrl, sgTENT5A-1, and sgTENT5A-2 following stimulation with CT-DNA (1 μg/mL) or infection with HSV-1 (MOI = 1) for the indicated time points. Data are presented as mean ± S.D. Two-way ANOVA; \**P <* 0.05, \*\**P <* 0.01, \*\*\**P <* 0.001, \*\*\*\**P <* 0.0001 (A). (B) Western blot analysis of ISG15 as well as phosphorylated and total STING, TBK1 and IRF3 in THP-1 cells stably expressing sgCtrl, sgTENT5A-1, and sgTENT5A-2 following stimulation with CT-DNA (1 μg/mL) for the indicated time. (C) Western blot analysis of ISG15, ICP0, as well as phosphorylated and total STING, TBK1 and IRF3 in THP-1 cells stably expressing sgCtrl, sgTENT5A-1, and sgTENT5A-2 following infection with HSV-1 (MOI = 1) for the indicated time.

**Supplementary Figure 7.**
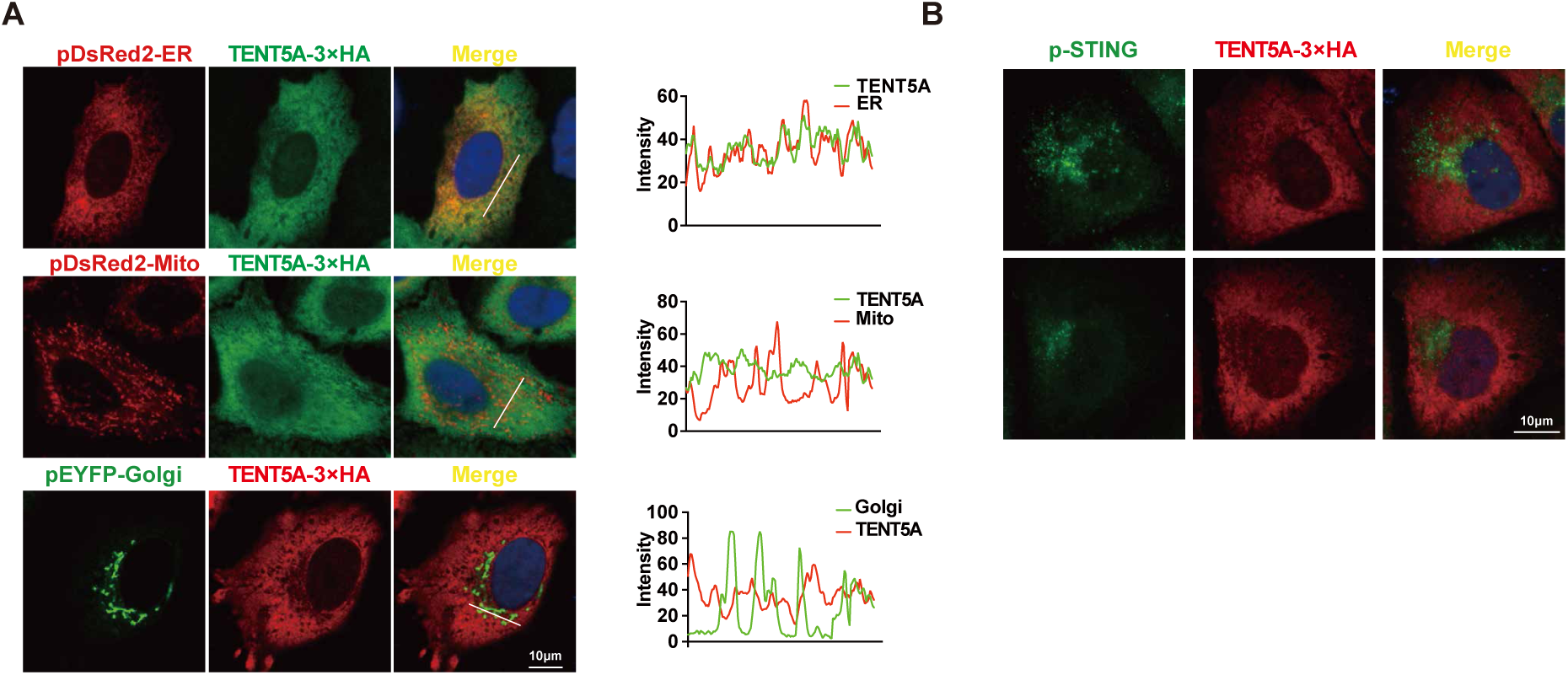
Subcellular localization of TENT5A. (A) Colocalization of TENT5A-3×HA with ER-Tracker (Red), Mito-Tracker (Red), or Golgi-Tracker (Green) in HeLa cells. The colocalization was quantified using ImageJ. Scale bar, 10 μm. (B) Colocalization of TENT5A-3×HA with endogenous p-STING in HeLa cells. The colocalization was quantified using ImageJ. Scale bar, 10 μm.

**Supplementary Figure 8.**
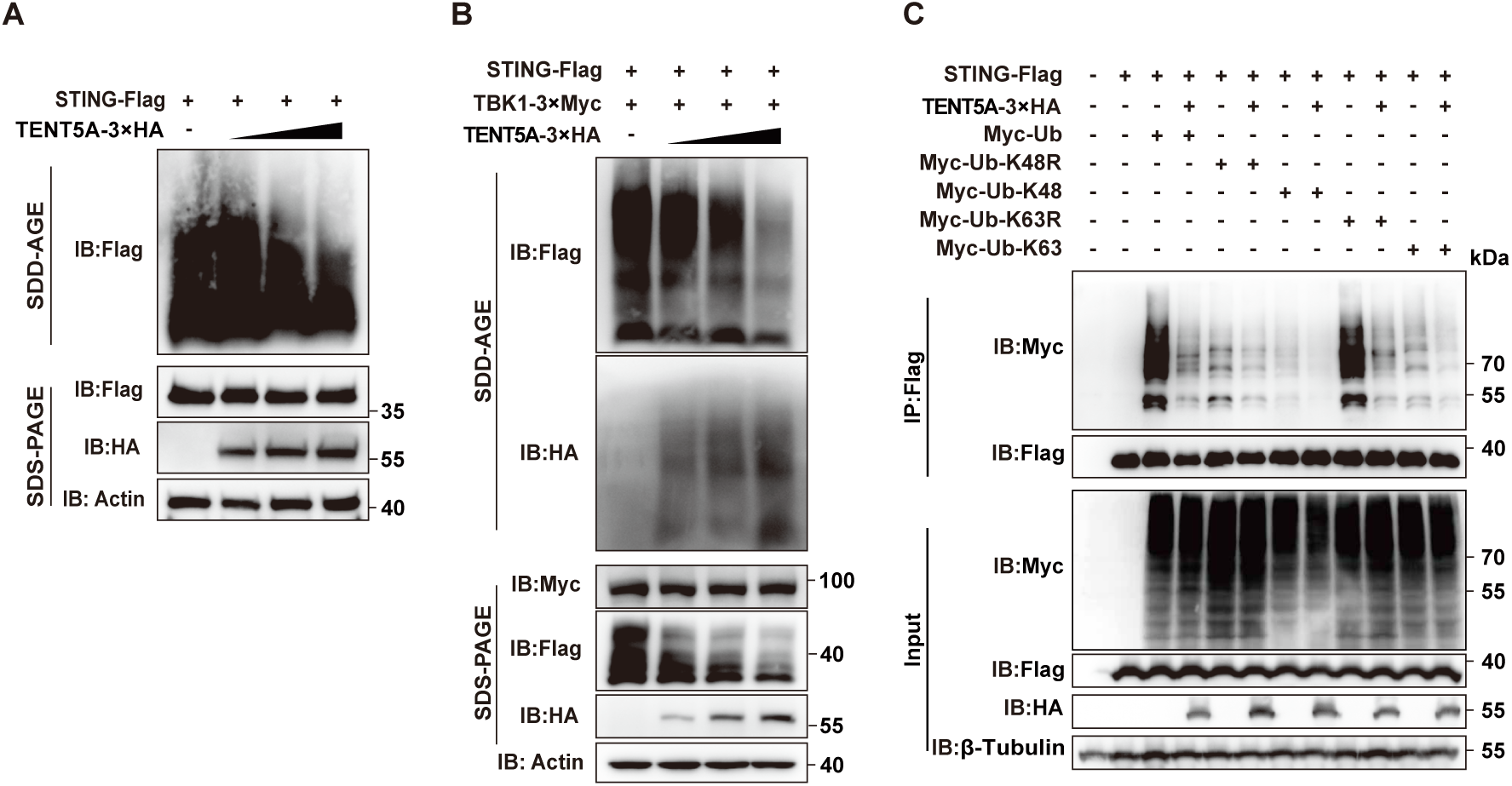
TENT5A inhibits STING oligomerization and K48, K63 Ubiquitination. (A) SDD-AGE analysis of TENT5A effect on STING oligomerization. HEK293T cells were co-transfected with STING-Flag and increasing doses of TENT5A-3×HA expression plasmids. (B) SDD-AGE analysis of TENT5A effect on STING oligomerization. HEK293T cells were co-transfected with STING-Flag, TBK1-3×Myc and increasing doses of TENT5A-3×HA expression plasmids. (C) Co-IP assays using anti-Flag beads to assess TENT5A effect on STING ubiquitination. HEK293T cells were co-transfected with STING-Flag, TENT5A-3×HA plasmids, and ubiquitin-related constructs (Myc-Ub, Myc-Ub-K48R, Myc-Ub-K48, Myc-Ub-K63R, Myc-Ub-K63).

**Supplementary Figure 9.**
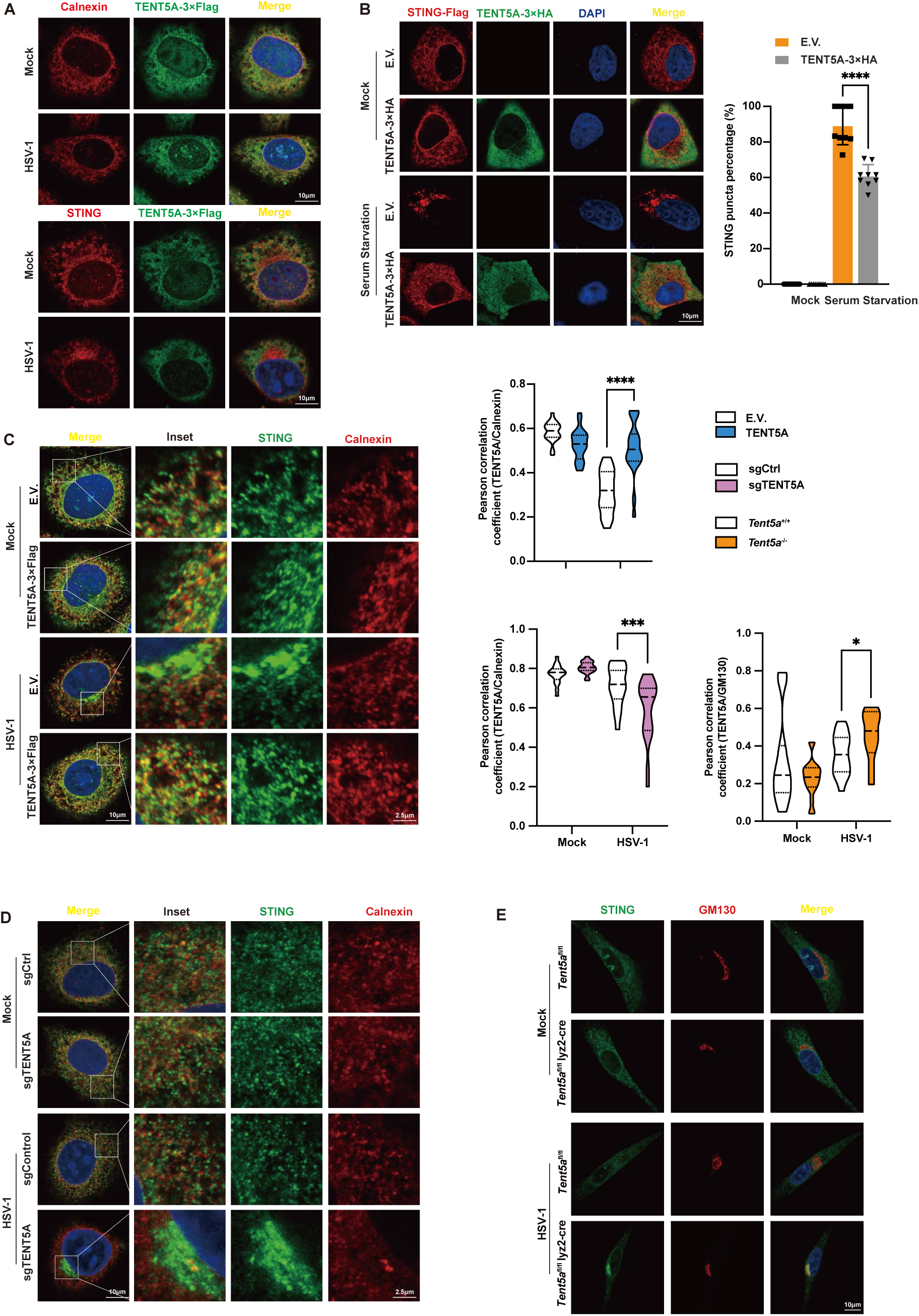
TENT5A inhibits STING ER-Golgi translocation. (A) Immunofluorescence colocalization analysis of TENT5A-3×Flag and endogenous Calnexin and STING in HeLa cells after HSV-1 (MOI = 1, 5 h) infection. Scale bar, 10 μm. (B) Immunofluorescence colocalization analysis of TENT5A-3×Flag and STING-Flag in HeLa cells. Scale bar, 10 μm. (C, D) HeLa cells stably expressing E.V./TENT5A-3×Flag (C) or sgCtrl/sgTENT5A (D) were mock-infected or infected with HSV-1 (MOI = 1, 5 h). Immunofluorescence staining was performed to to assess STING colocalization with the ER marker Calnexin. Colocalization was quantified by Pearson’s correlation coefficient in 20 cells using ImageJ software. Scale bar, 10 μm (main image), 2.5 μm (inset). (E) *Tent5a*^fl/fl^ and *Tent5a*^fl/fl^ Lyz2-Cre BMDMs were mock-infected or infected with HSV-1 (MOI = 1, 5 h). Immunofluorescence staining was performed to assess STING colocalization with Calnexin. Colocalization was quantified as in (C, D). Scale bar, 10 μm. Data are presented as mean ± S.D. Two-tailed unpaired Student’s t-test; \**P <* 0.05, \*\**P <* 0.01, \*\*\**P <* 0.001, \*\*\*\**P <* 0.0001 (B, C, D, E).

**Supplementary Figure 10.**
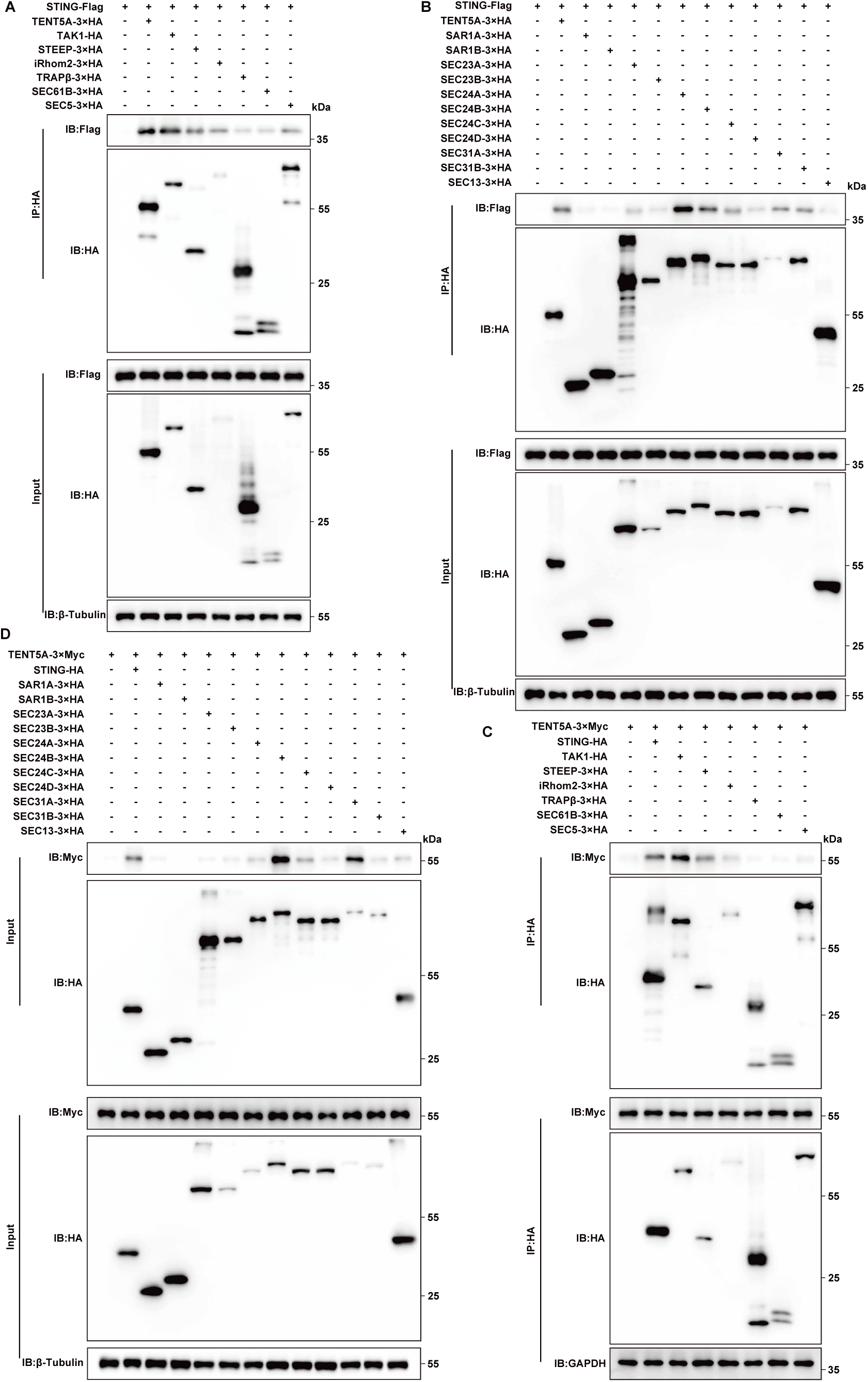
The interaction between STING translocation-related proteins with STING or TENT5A. (A, C) Co-IP assays using anti-HA beads to detect interactions of STING (A) or TENT5A (C) with STING trafficking regulators. HEK293T cells were co-transfected with STING-Flag (A) or TENT5A-3×Myc (C) and STING trafficking regulators constructs (TAK1-HA, STEEP-3×HA, iRhom2-3×HA, TRAPβ-3×HA, SEC61B-3×HA, SEC5-3×HA). (B, D) Co-IP assays using anti-HA beads to examine interactions of STING (B) or TENT5A (D) with COPII subunits. HEK293T cells were co-transfected with STING-Flag (B) or TENT5A-3×Myc (D) and COPII subunits constructs (SAR1A-3×HA, SAR1B-3×HA, SEC23A-3×HA, SEC23B-3×HA, SEC24A-3×HA, SEC24B-3×HA, SEC24C-3×HA, SEC24D-3×HA, SEC31A-3×HA, SEC31B-3×HA, SEC13-3×HA).

**Supplementary Figure 11.**
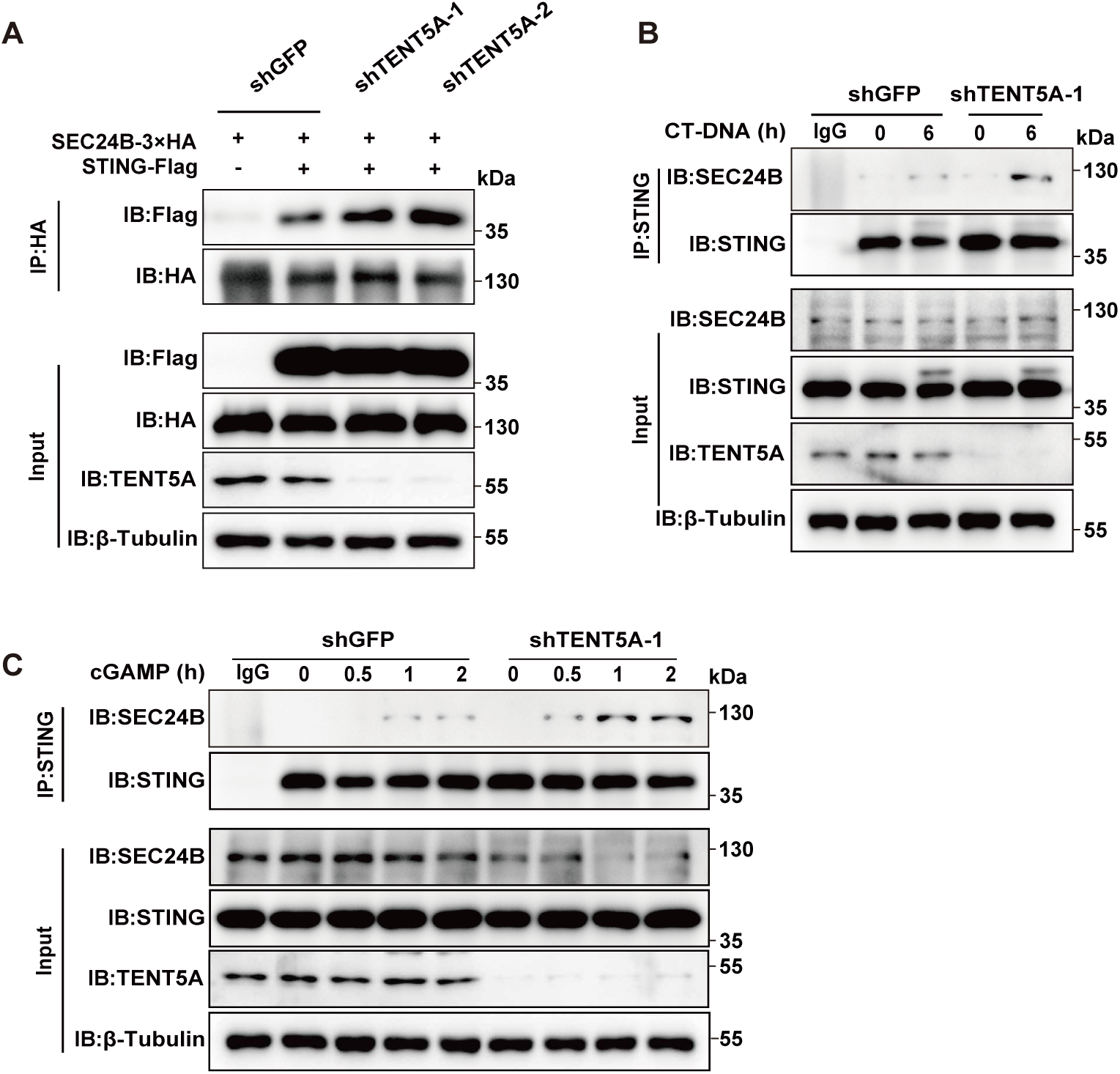
TENT5A knockdown promotes the interaction between STING and SEC24B. (A) Co-IP assays using anti-HA beads to assess the effect of TENT5A on the STING-SEC24B interaction. HEK293T cells stably expressing shGFP, shTENT5A-1, and shTENT5A-2 were co-transfected with STING-Flag and SEC24B-3×HA plasmids. (B) Co-IP assays using anti-STING antibody to detect the effect of TENT5A on the STING-SEC24B interaction. HeLa cells stably expressing shGFP or shTENT5A-1 were stimulated with CT-DNA (1 μg/mL, 6 h). (C) Co-IP assays using anti-STING antibody to analyze the effect of TENT5A on the STING-SEC24B interaction. THP-1 cells stably expressing shGFP or shTENT5A-1 were stimulated with cGAMP (1μg/mL) at the indicated time points.

**Supplementary Figure 12.**
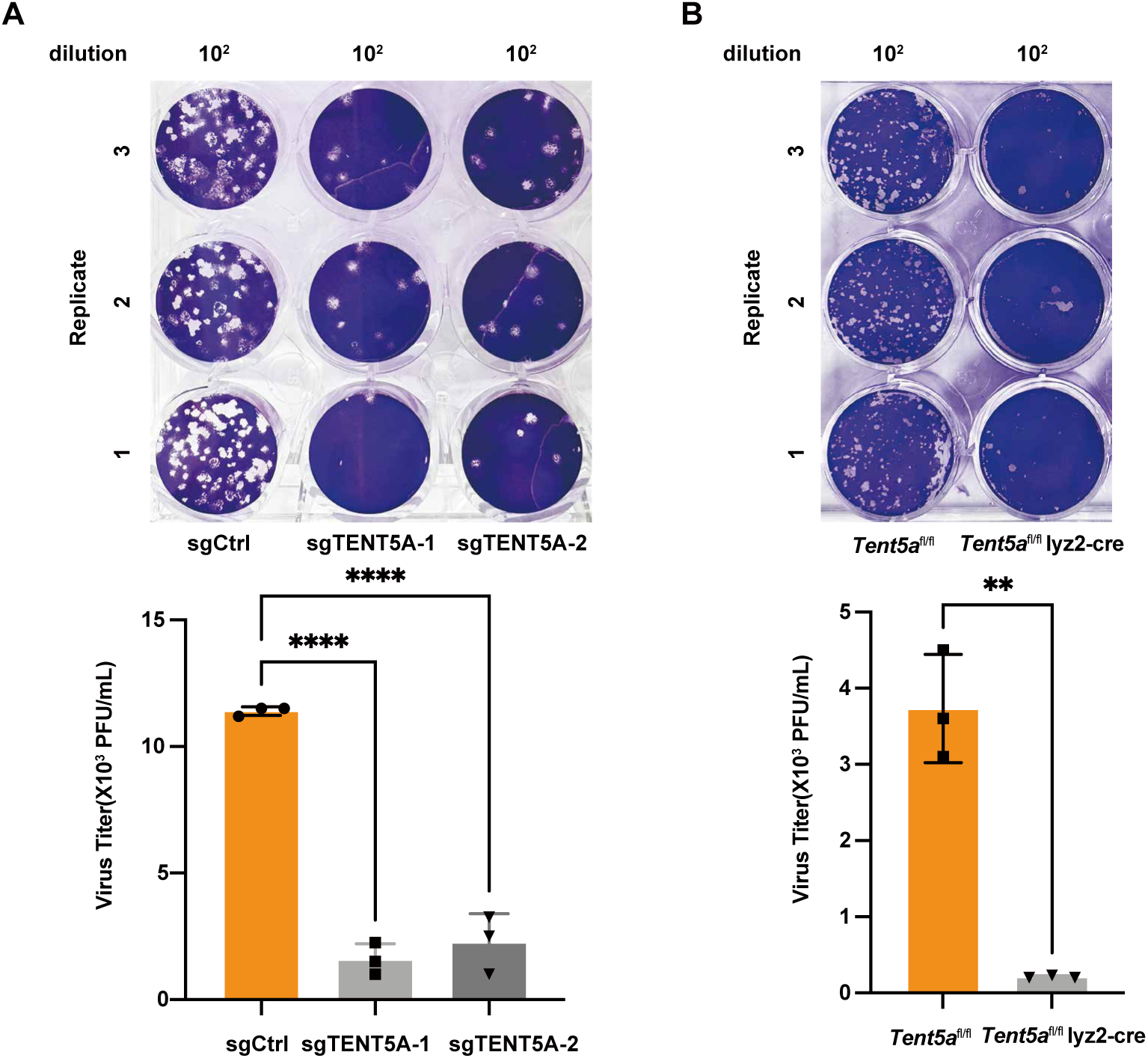
TENT5A promotes intracellular DNA virus replication. (A) HeLa cells with stable knockout for sgCtrl, sgTENT5A-1, and sgTENT5A-2 were infected with HSV–1 (MOI = 1, 24 h). Cell culture supernatants were harvested, and viral titers were determined by plaque assay. (B) BMDMs derived from T*ent5a*^fl/fl^ and *Tent5a*^fl/fl^ Lyz2-Cre mice were infected with HSV–1 (MOI = 1, 24 h). Cell culture supernatants were collected and viral titers were measured by plaque assay. Data are presented as mean ± S.D. Two-tailed unpaired Student’s t-test; \*\**P <* 0.01, \*\*\*\**P <* 0.0001.

## Discussion

The cGAS-STING pathway is a central innate immune signaling axis that detects cytosolic DNA and initiates type I interferon responses critical for antiviral and antitumor immunity. Since its discovery, extensive work has defined the core signaling cascade in which cGAS-derived cGAMP activates STING at the endoplasmic reticulum (ER), leading to TBK1 recruitment and IRF3 activation^[1, 2]^. However, it has become increasingly clear that STING activation is not solely a biochemical event but critically depends on spatial reorganization within the secretory pathway^[20, 21]^.

A major conceptual advance in recent years is the recognition that STING signaling is tightly coupled to its intracellular trafficking. STING must exit the ER and traffic to the ER–Golgi intermediate compartment and Golgi apparatus to become fully signaling competent^[5, 6]^. This process is now understood to be mediated by COPII vesicle formation machinery, with SEC24B functioning as a key cargo adaptor required for STING ER export^[22, 23]^. More recent structural and cell biological studies further suggest that STING trafficking is actively regulated by post-translational modifications and vesicular sorting signals, highlighting that ER exit is a rate-limiting and regulated step rather than a constitutive process^[8]^.

Despite these advances, current models of STING regulation remain largely centered on two mechanisms: (i) modulation of STING activation through post-translational modifications such as ubiquitination, phosphorylation, and palmitoylation^[12, 16, 24]^, and (ii) ER retention mediated by scaffold proteins such as STIM1 that prevent premature activation under basal conditions^[25]^. In both paradigms, STING trafficking is generally considered a downstream consequence of activation or sequestration dynamics rather than a process actively gated at the level of COPII cargo selection.

Here, our study identifies TENT5A as a previously unrecognized ER-resident checkpoint that directly controls STING access to the COPII trafficking machinery. Unlike STIM1, which restrains STING through physical sequestration at the ER membrane^[25]^, or ubiquitin-mediated pathways that regulate STING stability and signaling output^[14, 15]^, TENT5A operates upstream by competitively interfering with SEC24B binding to STING. This establishes a mechanistically distinct regulatory principle in which immune signaling competency is determined at the level of vesicular cargo selection.

Importantly, this finding revises the prevailing view of COPII machinery in innate immunity. SEC24B has been characterized as a positive regulator of STING ER export^[26]^, suggesting that COPII-mediated trafficking is primarily permissive. However, our data support a revised model in which COPII-dependent cargo loading is not constitutive but actively constrained by ER-resident checkpoint proteins. This introduces a competitive regulatory layer in which signaling substrates and trafficking adaptors directly compete for access to vesicular machinery, thereby integrating immune signaling control with secretory pathway organization.

Recent studies have increasingly emphasized the ER as an active regulatory hub in innate immunity rather than a passive reservoir for STING localization. Within this emerging framework, our findings define a spatial licensing mechanism in which STING activation is contingent upon successful engagement with COPII machinery. TENT5A imposes a threshold at this licensing step, thereby preventing inappropriate activation under homeostatic conditions while permitting robust signaling upon pathogen detection. This expands current models of innate immunity from biochemical activation thresholds to include spatially defined checkpoints at organelle boundaries. Functionally, disruption of this checkpoint enhances type I interferon responses, improves antiviral defense against DNA viruses, and promotes antitumor immunity in vivo, underscoring the physiological importance of spatial regulation in immune homeostasis. These observations suggest that modulation of ER trafficking checkpoints may represent a previously underexplored therapeutic strategy for infectious disease, cancer, and autoinflammatory disorders.

In summary, we identify TENT5A as a competitive ER checkpoint that regulates STING activation by controlling its access to COPII-dependent trafficking machinery. This work revises current models of STING regulation by introducing vesicular cargo selection as an active immune regulatory node and establishes spatial licensing at the ER as a fundamental principle governing innate immune signaling.

## Material and Methods

### Cells and Viruses

HEK293T, HeLa, THP-1, Vero and B16-Luc cells were obtained from the American Type Culture Collection (ATCC). HEK293T and Vero cells were cultured in Dulbecco’s modified Eagle’s medium (DMEM, Gibco) supplemented with 10% fetal bovine serum (FBS, Gibco) and 1% penicillin-streptomycin (Gibco), while B16-Luc and THP-1 cells were grown in RPMI 1640 medium (Gibco) supplemented with 10% FBS and 1% penicillin-streptomycin. HeLa cells were cultured in DMEM supplemented with 10% FBS and 1% penicillin-streptomycin under routine conditions and shifted to DMEM with 1% FBS for 48 h prior to dsDNA transfection or DNA virus infection, as reported earlier^[27]^. THP-1 cells were differentiated into macrophages by overnight treatment with 150 nM PMA. After removal of PMA, cells were incubated for an additional 24 h prior to experiments. All cells were maintained at 37 °C in a humidified atmosphere with 5% CO₂ and 95% relative humidity.

Herpes simplex virus 1 (HSV-1) was kindly provided by Prof. Chengjiang Gao (Shandong University, China), and vaccinia virus (VACV) was a gift from Prof. Zhengfan Jiang (Peking University, China).

### Isolation of mouse embryonic fibroblasts (MEFs), peritoneal macrophages (PMs) and bone-marrow-derived macrophages (BMDMs)

MEFs, PMs and BMDMs have been described elsewhere^[28]^. Briefly, MEFs were harvested from E15 embryos of *Tent5a*^+/+^ and *Tent5a*^-/-^ mice. For PMs isolation, *Tent5a*^fl/fl^ and *Tent5a*^fl/fl^ Lyz2-Cre mice received an intraperitoneal (i.p.) injection of BD Bacto™ Thioglycollate Broth, and PMs were collected 4 days later. Both MEFs and PMs were cultured in DMEM supplemented with 10% fetal bovine serum (FBS). BMDMs were isolated from the tibias and femurs of *Tent5a*^fl/fl^ and *Tent5a*^fl/fl^ Lyz2-Cre mice, and differentiated for 7 days in DMEM containing 20% FBS, 2 mM L-glutamine and 30% L929-conditioned supernatant.

### Sequences, Plasmids and Transfection

cDNAs of human TENT5A, STEEP, iRhom2, TRAPβ, SEC61B, SEC5, SAR1A, SAR1B, SEC23A, SEC23B, SEC24A, SEC24B, SEC24C, SEC24D, SEC31A, SEC31B and SEC13 were amplified from THP-1 cells via conventional PCR and subcloned into the pXJ2 vector. All other plasmids applied in this work have been described elsewhere^[29–31]^.

For stable cell line construction, overexpression constructs were generated with the pLVX-IRES-Puro vector. Short hairpin RNAs (shRNAs) and single-guide RNAs (sgRNAs) were separately cloned into the pLKO.1-Puro and pLenti CRISPRv2-Puro vectors, respectively. Detailed sequences of TENT5A-targeting shRNAs and sgRNAs are provided in Table S1. Plasmids were transfected into cells using Lipofectamine 3000 (Invitrogen), and CT-DNA or cGAMP was transfected with Lipofectamine 2000 (Invitrogen).

### Antibodies and Reagents

Primary Antibodies

The primary antibodies applied in this work are listed below: anti-HA-Tag (TU281749L, M20003H; Abmart), anti-MYC-Tag (16286-1-AP; Proteintech), anti-FLAG-Tag (600-401-383; Rockland, M20008H; Abmart, ab213519; Abcam), anti-Lamin B1 (AB0054; Abways), anti-β-Tubulin (M30109H; Abmart), anti-GAPDH (M20006H; Abmart), anti-Actin (M20011H; Abmart), anti-Calnexin (66903-1-Ig; Proteintech), anti-GM130 (66662-1-Ig; Proteintech), anti-HSV1 ICP0 (ab6513; Abcam), anti-ISG15 (CY7086; Abways), anti-p-STING (Ser366/Ser365, 19781/72971; CST), anti-p-IRF3 (CY6575; Abways), anti-p-TBK1 (Ser172, 5483; CST), anti-IRF3 (CY5779; Abways), anti-TBK1 (CY5145; Abways), anti-STING (13647; CST, 19851-1-AP/66680-1-Ig; Proteintech), anti-TENT5A (PA5-23898/PA5-120184; Thermo), anti-SEC24B (PC4452M; Abmart), anti-TMP21 (15199-1-AP; Proteintech).

### Flow Cytometry Antibodies

PE-conjugated CD69, APC-conjugated CD62L, FITC-conjugated CD45 and Pacific Blue-conjugated CD8a (104507/104411/147710/100728; BioLegend) were used for flow cytometric analysis.

Immunoprecipitation (IP) Reagents

Protein A/G agarose and magnetic beads (A10001/A10002; Abmart), anti-FLAG magnetic beads, and anti-Myc/HA agarose beads (M20118L/M20012L/M20013L; Abmart).

Chemicals and Small Molecules

Calf thymus DNA (CT-DNA, D8020; Solarbio), 2′3′-cGAMP (9523-46-01; InvivoGen), D-luciferin sodium salt (D9390; Solarbio), Digitonin (D806779; Macklin), Creatine phosphate (C916918; Macklin), Creatine kinase (C902501; Macklin), Guanosine triphosphate (GTP, G810427; Macklin) and Adenosine triphosphate (ATP, A8270; Solarbio).

### Mice

*Tent5a* conditional knockout (*Tent5a*^fl/-^) and germline heterozygous knockout (*Tent5a*^+/-^) mice were purchased from GemPharmatech (Nanjing, China). Lyz2-Cre transgenic mice were obtained from Cyagen Biosciences (Suzhou, China). All experimental mice were bred on a C57BL/6J background and kept in individually ventilated cages (IVC) within the Specific Pathogen-Free (SPF) animal facility of the Laboratory Animal Center, Shandong University.

Animal experiments were conducted using male and female mice aged 8 to 12 weeks. All procedures complied with the guidelines of the Association for Assessment and Accreditation of Laboratory Animal Care, and were approved by the Institutional Animal Care and Use Committee (IACUC) of Shandong University.

### Viral Infection and Plaque Assay

For viral infection, HeLa, Vero, THP-1 cells, BMDMs, MEFs and PMs were seeded 24 h in advance. On the day of infection, cells were rinsed with serum-free medium and incubated with HSV-1 or VACV (diluted in serum-free medium) at the indicated multiplicity of infection (MOI) for 1 h to allow viral adsorption. The inoculum was then discarded, and cells were cultured in fresh complete medium.

Viral titers were quantified by plaque assay using Vero cells. Briefly, Vero cells were seeded into 24-well plates 24 h before use. Cells were exposed to serially diluted HSV-1 supernatants for 30 min, followed by addition of DMEM supplemented with 2% FBS and 0.5% low-melt agarose. Once the overlay solidified, cells were incubated at 37 °C for 48-72 h. For fixation, cells were treated with a 1:1 methanol-ethanol mixture at 4 °C for 6 h. After removal of the agarose layer, cells were stained with 0.05% crystal violet solution. Plates were rinsed with distilled water and air-dried. Viral plaques were counted manually, and viral titers were calculated accordingly.

### mRNA Isolation and Quantitative Reverse Transcription PCR

Total cellular RNA was isolated from cells with TRIzol reagent (Invitrogen). Purified RNA was reverse transcribed into complementary DNA (cDNA) using the HiScript III 1st Strand cDNA Synthesis Kit (Vazyme). Quantitative reverse transcription real-time PCR (RT-qPCR) was performed on a Roche LightCycler 96 platform with UltraSYBR Mixture (CWBIO), using appropriately diluted cDNA as the template.

All experiments were conducted following the manufacturer’s instructions. Primer sequences for target genes are provided in Table S2. The relative expression of target genes was calculated by the 2^−ΔΔCt^method, with GAPDH serving as the internal reference gene.

### Lentivirus Production and Infection

For lentivirus packaging, HEK293T cells were co-transfected with transfer plasmids and lentiviral packaging plasmids pMD2.G and psPAX2 using Lipofectamine 2000. The transfer plasmids included TENT5A overexpression constructs (pLVX-IRES-Puro), TENT5A-targeting shRNA constructs (pLKO.1-Puro), TENT5A-targeting sgRNA constructs (pLenti CRISPRv2-Puro), and the matched empty vector controls. Culture supernatants containing lentiviral particles were collected at 48 h post-transfection.

HeLa and THP-1 cells were incubated with the corresponding lentiviruses in medium containing 8 μg/mL Polybrene for three consecutive days to establish stable cell lines. Afterwards, cells were cultured in medium supplemented with puromycin for 7 days to screen stable cell lines. The efficiency of gene overexpression, knockdown and CRISPR knockout was verified via Western blotting at 1-week post-transduction.

### Luciferase Reporter Assay

Dual-luciferase reporter assays were performed as described previously^[30, 31]^. In brief, HEK293T cells were co-transfected with firefly luciferase reporter plasmids (pGL3-IFN-β-Luc, pGL3-IFN-λ1-Luc and pGL3-ISRE-Luc) and constructs expressing distinct signaling molecules, in the presence or absence of the TENT5A overexpression plasmid, as indicated in the figure legends. The Renilla luciferase reporter plasmid pRL-TK was included in each transfection as an internal control to calibrate transfection efficiency. Cells were harvested and lysed at 36 h post-transfection. Luciferase activity was detected using the Dual-Luciferase Reporter Assay Kit (Vazyme) following the manufacturer’s instructions. Relative luciferase activity was presented as the ratio of firefly luciferase activity to Renilla luciferase activity.

### Co-immunoprecipitation and immunoblotting

Co-IP assays were performed as described previously^[28]^. Briefly, transfected HEK293T cells were lysed with pre-chilled lysis buffer (50 mM Tris-HCl, pH 7.4, 50 mM EDTA, 1.0% NP-40, 150 mM NaCl) containing protease and phosphatase inhibitors (Sigma). Lysates were rotated at 4 °C for 30 min and centrifuged at 15,000 × g for 20 min. The resulting supernatants were harvested, and total protein concentrations were determined via the BCA assay (Thermo). Samples were incubated with target-specific antibodies together with Protein A/G agarose beads (Thermo), or directly incubated with anti-Flag, anti-HA or anti-Myc affinity beads (Thermo/Abmart) overnight at 4 °C. After four rounds of washing with lysis buffer (5 min for each wash), the bead-bound protein complexes were eluted by boiling in 2× SDS loading buffer for immunoblot analysis.

For immunoblotting, whole-cell lysates were denatured in 5× SDS loading buffer at 100 °C for 10 min. Cytoplasmic and nuclear protein fractions were isolated using a commercial extraction kit (Beyotime). Proteins were resolved by SDS-PAGE and electrotransferred to PVDF membranes (Millipore). Membranes were blocked with 5% non-fat milk in TBST buffer. Subsequently, membranes were probed with primary antibodies at 4 °C overnight, and then incubated with corresponding secondary antibodies at room temperature for 1 h. Protein signals were detected using an ECL chemiluminescence system (Invitrogen).

### Immunofluorescence microscopy

HeLa cells and BMDMs were seeded onto poly-L-lysine-coated glass coverslips (Sigma) in 12-well culture plates. After 24 h, cells were subjected to plasmid transfection, viral infection, or stimulation with CT-DNA or cGAMP for the indicated time points. For immunofluorescence staining, cells were rinsed thoroughly with phosphate-buffered saline (PBS) and fixed with pre-chilled methanol at −20°C for 15 min. Cell permeabilization and blocking were performed using a commercial immunofluorescence assay kit (Beyotime) in accordance with the manufacturer’s instructions. Samples were incubated with primary antibodies at 4 °C overnight. After three washes with PBS, cells were incubated with fluorophore-conjugated secondary antibodies at room temperature for 1 h. Coverslips were mounted with DAPI-containing anti-fade mounting medium (Beyotime). Fluorescence images were acquired with a Zeiss LSM900 laser scanning confocal microscope, and quantitative analysis was carried out using ImageJ software.

### Flow Cytometry

Flow cytometric analysis was performed to characterize CD8⁺ T cell subsets in melanoma tissues using a Beckman Coulter Gallios flow cytometer. Single-cell suspensions were isolated from harvested melanoma tissues. Cell surface staining was conducted by incubating cells with fluorophore-conjugated antibodies targeting CD45, CD8, CD69 and CD62L at room temperature for 30 min.After incubation, cells were washed twice with phosphate-buffered saline (PBS) to eliminate unbound antibodies. The final cell suspensions were resuspended in PBS and immediately analyzed by flow cytometry.

### In vivo HSV-1 infection model

Age- and sex-matched *Tent5a*^fl/fl^ and *Tent5a*^fl/fl^ Lyz2-Cre mice were inoculated with HSV-1 via the tail vein at a dose of 1 × 10⁷ PFU per mouse. At 72 h post-infection, lung, spleen, liver, brainstem, and peripheral blood were collected for qRT-PCR detection. Serum cytokine concentrations were determined by ELISA, and HSV-1 viral loads in Liver tissues were quantified via plaque assay using Vero cells.

For survival experiments, mice received HSV-1 by intravenously administered at 1 × 10⁸ PFU per mouse. Animal survival was monitored daily for 7 days. Lung tissues isolated from naive and HSV-1-infected mice were fixed in 10% neutral buffered formalin, followed by paraffin embedding, tissue sectioning and hematoxylin and eosin (H&E) staining. Histopathological lesions were examined and evaluated using a light microscope.

### Semi-denaturing detergent agarose gel electrophoresis (SDD–AGE)

SDD-AGE was conducted following a previously reported protocol with slight modifications^[28, 32]^. Briefly, cells were lysed using pre-chilled lysis buffer (50 mM Tris-HCl, pH 7.4, 150 mM NaCl, 1.0% NP-40, 5 mM EDTA) containing freshly prepared protease and phosphatase inhibitor cocktails. Cell lysates were incubated on ice for 30 min, then centrifuged at 12,000 × g for 15 min at 4 °C to pellet cell debris. The resulting supernatants containing native STING protein were harvested for electrophoresis.

Protein samples were mixed with 1× SDD-AGE loading buffer (0.5× TBE, 10% glycerol, 2% SDS, 0.0025% bromophenol blue). To preserve the oligomeric state of STING, samples were loaded onto a 1.5% agarose gel without heat denaturation. Electrophoresis was run in 1× TBE buffer supplemented with 0.1% SDS at a constant voltage of 100 V for 35 min at 4 °C. Following electrophoresis, STING oligomers were electrotransferred to Immobilon PVDF membranes (Millipore) and detected by immunoblotting.

### Native polyacrylamide gel electrophoresis (PAGE)

Native PAGE was utilized to assess the oligomeric status of STING. This assay was modified from a classic protocol widely used for detecting IRF3 dimerization^[32]^. Before sample loading, gels were pre-run at a constant current of 45 mA for 30 min with running buffer (25 mM Tris-Cl, pH 8.4, 192 mM glycine). Specifically, the cathode buffer was supplemented with 0.2% deoxycholate, while the anode buffer contained no deoxycholate.Treated cells were lysed with ice-cold IP lysis buffer containing freshly prepared protease and phosphatase inhibitor cocktails. Cell lysates were centrifuged at 10,000 × g for 10 min at 4 °C to remove cell debris. Clarified supernatants were mixed with native loading buffer (125 mM Tris-Cl, pH 6.8, 30% glycerol, 0.002% bromophenol blue). Electrophoresis was carried out at 25 mA constant current for 120 min in an ice bath to maintain the native structural conformation of STING and prevent protein unfolding.

### Tumor models

Age- and sex-matched *Tent5a*^fl/fl^ and *Tent5a*^fl/fl^ Lyz2-Cre mice were used for tumor implantation. B16-Luc cells (5 × 10⁵) were resuspended in 100 μL phosphate-buffered saline (PBS) and injected subcutaneously (s.c.) into the right dorsal flank of each mouse. For in vivo bioluminescence imaging, mice received an intraperitoneal (i.p.) injection of D-luciferin sodium salt (Solarbio) at a dose of 150 mg/kg on days 8 and 14 post tumor inoculation. Bioluminescence signals were detected with a ClinX IVScope 8500 Pro in vivo imaging system.

### Enzyme-linked immunosorbent assay (ELISA)

The levels of IFN-β in cell culture supernatants and mouse sera were quantified using a commercial ELISA kit (Multi Science), following the manufacturer’s protocols.

### Isolation of ER and Golgi fractions

ER and Golgi apparatus fractions were isolated using a modified version of a well-established protocol^[33]^. In brief, 1.5 × 10⁷ HeLa cells were harvested and centrifuged at 1000 × g for 5 min. Cell pellets were washed three times with pre-chilled 1× PBS, then resuspended in hypotonic buffer (20 mM HEPES-KOH, pH 7.2, 10 mM KCl, 3 mM MgCl₂) supplemented with freshly prepared protease and phosphatase inhibitor cocktail. Cells were homogenized by repeated passage through a 25-gauge syringe needle for 25 times. Cell homogenates were subjected to sequential centrifugation at 1,000 × g and 12,000 × g for 10 min each to remove intact cells, cell debris, nuclei and mitochondria. The clarified supernatant was mixed with 60% sucrose solution to reach a final concentration of 37% (w/w). A discontinuous sucrose density gradient was layered from bottom to top: 0.5 mL of 42% (w/w) sucrose cushion, 1.5 mL of 37% (w/w) sample solution, 0.5 mL of 35% (w/w) sucrose, 0.5 mL of 29% (w/w) sucrose, and 1 mL of 20% (w/w) sucrose as the top layer. All gradient solutions were prepared using gradient buffer (10 mM HEPES-KOH, pH 7.4, 0.25 M sucrose, 1 mM EDTA) and added with fresh protease and phosphatase inhibitors. Samples were loaded into centrifuge tubes (331379, Beckman), and the remaining void volume was filled with mineral oil. Ultracentrifugation was carried out at 150,000 × g for 3 h at 4 °C with an SW41 Ti rotor. After centrifugation, 200 μL aliquots of gradient fractions were collected. Proteins from each fraction were concentrated by methanol-chloroform precipitation and analyzed via immunoblotting.

### In Vitro COPII Budding Assay

Preparation of cytosolic fractions: Cytosolic extracts were prepared according to well-documented protocols^[33, 34]^. In brief, 1.5 × 10⁷ HeLa cells were harvested and resuspended in pre-chilled HEPES buffer (20 mM HEPES, pH 7.3, 110 mM potassium acetate, 2 mM magnesium acetate) containing freshly prepared protease and phosphatase inhibitor cocktail as well as 0.3 mM dithiothreitol (DTT). Cells were homogenized by repeated passage through a 22-gauge syringe needle for 60 times. The homogenate was ultracentrifuged at 160,000 × g for 2 h to remove insoluble material, and the resulting supernatant was collected for downstream experiments.

Preparation of semi-permeabilized cells: Semi-permeabilized HeLa cells were generated using a modified standard protocol^[23, 35]^. A total of 1.5 × 10⁷ HeLa cells were harvested and resuspended in 10 mL pre-chilled HEPES buffer. After centrifugation at 250 × g for 3 min at 4 °C, the supernatant was discarded, and cell pellets were resuspended in 6 mL KHM buffer (50 mM HEPES, pH 7.3, 90 mM potassium acetate). Cells were incubated with 10 μg/mL digitonin on ice for 5 min to permeabilize the plasma membrane. The reaction was stopped by adding 8 mL KHM buffer, and cells were collected by centrifugation at 250 × g for 3 min at 4 °C. Cell pellets were resuspended in 10 mL HEPES buffer, kept on ice for 10 min, and centrifuged again under the same conditions. The final pellets were resuspended in 1 mL KHM-Cl buffer (110 mM KCl, 20 mM HEPES, pH 7.3, 2 mM MgCl₂) and centrifuged at 10,000 × g for 15 s at 4 °C. Cells were diluted with KHM-Cl buffer and adjusted to the appropriate OD₆₀₀, then used immediately for the COPII budding assay.

In vitro COPII vesicle reconstitution assay: A modified cell-free system was used to reconstitute COPII vesicle budding and STING trafficking^[35, 36]^. The 50 μL reaction mixture contained cytosol (2 mg/mL) and semi-permeabilized cells (OD₆₀₀ = 10) diluted in KHM-Cl buffer. An energy regeneration system (40 mM creatine phosphate, 0.2 mg/mL creatine phosphokinase, 1 mM ATP and 0.2 mM GTP) and freshly prepared protease and phosphatase inhibitors were added to the reaction system. The mixture was incubated at 30 °C for 45 min to allow COPII vesicle formation, then placed on ice to terminate the reaction. For total input control, 10 μL of the reaction mixture was mixed with 40 μL 2× SDS loading buffer. The remaining supernatant was centrifuged at 20,000 × g for 20 min at 4 °C to remove cell debris, followed by ultracentrifugation at 100,000 × g for 1 h at 4 °C to pellet purified COPII vesicles. The vesicle pellets were resuspended in 30 μL 1× SDS loading buffer. All samples were boiled at 100 °C for 10 min and analyzed by immunoblotting.

### Statistical Analysis

All data are expressed as mean ± SD from no fewer than three independent experiments. The two-tailed Student’s t-test was used for comparisons between two groups. For multiple group analyses, one-way analysis of variance (ANOVA) combined with Tukey’s post-hoc multiple comparison test was applied. Statistical significance was denoted as \**P* < 0.05, \*\**P* < 0.01, \*\*\**P* < 0.001 and \*\*\*\**P* < 0.0001. All statistical analyses were performed using GraphPad Prism.

## Supplementary Tables

**Table S1.**
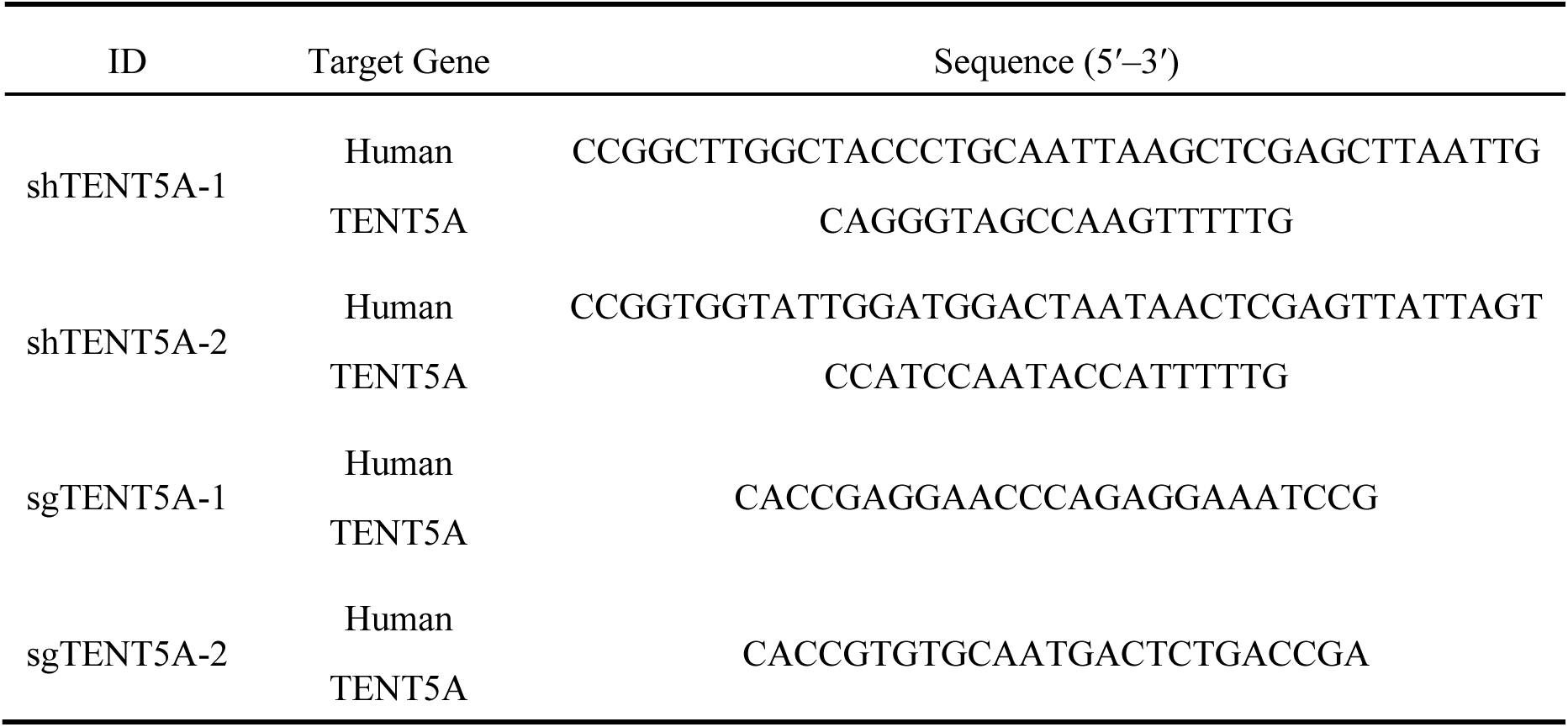
Information of shRNA and sgRNA sequences used in this study.

**Table S2.**
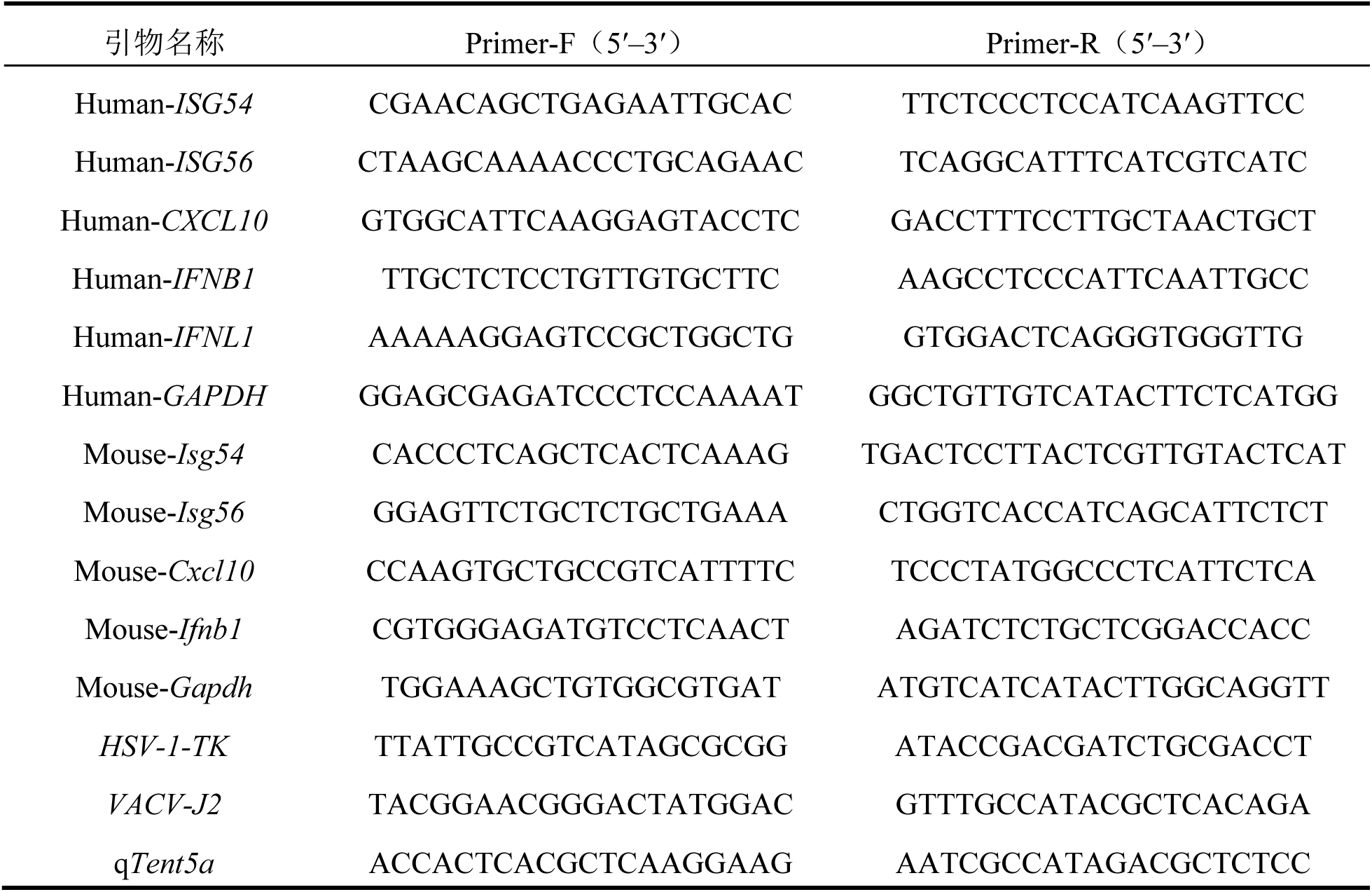
Primers used for quantitative real-time PCR.

